# Comparative Proteomic Analysis of Paired Human Milk Fat Globules and Membranes and Mouse Milk Fat Globules Identifies Core Cellular Systems Contributing to Mammary Lipid Trafficking and Secretion

**DOI:** 10.1101/2023.02.28.530322

**Authors:** Jayne F. Martin Carli, Jenifer Monks, James L. McManaman

## Abstract

Human milk delivers critical nutritional and immunological support to the infant. The milk fat globule and its membrane contain many bioactive components, yet the mechanism of milk fat secretion and how milk fat globule (MFG) components are regulated are poorly defined. In this study, we perform quantitative proteomic profiling of milk fat globules from human and mouse milk, as well as from isolated membranes physically disrupted from human milk fat globules.

Using paired analyses of the human samples we report membrane enrichment of the proteins involved in docking/tethering the lipid droplet to the membrane as well as minor components involved in the signaling pathway for secretion. Comparing abundance between human and mouse milk fat globules we find that 8 of 12 major milk fat globule proteins are shared between the two species. Comparative pathway enrichment analyses between human and mouse samples reveal similarities in shared membrane trafficking and signaling pathways involved in milk fat secretion. Our results advance knowledge of the composition and relative quantities of proteins in human and mouse milk fat globules in greater detail, provide a quantitative profile of specifically enriched human milk fat globule membrane proteins, and identify core cellular systems involved in milk lipid secretion.

## Introduction

Human milk is well appreciated to be the gold standard of nutrition for human infants (Victora, Bahl et al. 2016). Currently, there is rapidly expanding interest in more fully elucidating how the interactions between individual milk components make human milk greater than the sum of its parts (Christian, Smith et al. 2021). The broader “matrix” in which these individual milk components exist is thought to contribute to both infant and maternal health. Many factors are expected to affect the milk matrix, including behavioral, environmental, structural, and organizational influences. One of the most structurally and organizationally complex components of milk is the milk fat globule (MFG), suggesting that the MFG is a major contributor to the effects of the milk matrix. MFGs form a specialized lipid delivery system, providing nutritional components to support infant growth and development as well as immunological protection against disease. Their surrounding phospholipid trilayer membrane (milk fat globule membrane; MFGM) is unique to milk, as infant formulas have traditionally provided lipids in the form of vegetable oil. Recent efforts have focused on adding MFGMs to infant formulas in recognition of the immunological and neurodevelopmental benefits associated with the protein, carbohydrate, and lipid constituents of MFGMs (Brink and Lonnerdal 2020).

MFG production begins with triacylglycerol (TG) synthesis in the endoplasmic reticulum (ER) (Stein and Stein 1967, Kassan, Herms et al. 2013). Accumulating TGs are released into the cytoplasm where they are surrounded by an ER-derived phospholipid monolayer containing numerous attached or embedded proteins, resulting in formation of organelle structures referred to as cytoplasmic lipid droplets (CLD) (Dylewski, Dapper et al. 1984, Walther and Jr. 2012).

CLDs in mammary epithelial cells are coated with perilipin 2 (Plin2), which is thought to confer stability by protecting CLDs from lipolysis (Listenberger, Ostermeyer-Fay et al. 2007, Russell, Schaack et al. 2011). CLDs can fuse to form larger CLDs in process that is thought to be facilitated by cell death-inducing DNA fragmentation factor, alpha subunit-like effector A (Cidea), while concurrently trafficking toward the apical surface (Wooding 1971, Stemberger and Patton 1981, Wang, Lv et al. 2012, Wu, Zhou et al. 2014, Barneda, Planas-Iglesias et al. 2015, Monks, Dzieciatkowska et al. 2016, Masedunskas, Chen et al. 2017) of the polarized luminal mammary epithelial cell (MEC). While moving intracellularly, CLDs interact with other organelles, including the Golgi apparatus, mitochondria and casein-containing secretory vesicles (Wooding 1971, Wooding 1973, Stemberger, Walsh et al. 1984, Wu, Howell et al. 2000, Mather, Jack et al. 2001, Honvo-Houéto, Henry et al. 2016). When they arrive at the apical cytoplasm, CLDs form contacts with the apical plasma membrane via interactions between CLD-coating Plin2, cytoplasmic xanthine dehydrogenase (Xdh; also known as xanthine oxidoreductase, Xor) and the transmembrane apical plasma membrane protein butyrophilin, subfamily 1, member A1 (Btn1a1) (Ishii, Aoki et al. 1995, Keenan and Patton 1995, Mather and Keenan 1998, McManaman, Palmer et al. 2002, Vorbach, Scriven et al. 2002, Ogg, Weldon et al. 2004, Robenek, Hofnagel et al. 2006, Jeong, Rao et al. 2009, Monks, Dzieciatkowska et al. 2016, Jeong, Kadegowda et al. 2021). These proteins and their interactions allow for tight tethering, or docking, of CLDs to the apical plasma membrane, where they can continue to grow by fusion and protrude into the alveolar lumen (Dylewski, Dapper et al. 1984, Masedunskas, Chen et al. 2017). Oxytocin release from the pituitary, which is driven by nipple stimulation, drives contraction of the surrounding myoepithelial cells, causing CLDs to become enveloped by the apical plasma membrane bilayer and secreted into the alveolar lumen as MFGs by an apocrine mechanism (Kurosumi, Kobayashi et al. 1968, Mather and Keenan 1998, Mather, Jack et al. 2001, Masedunskas, Chen et al. 2017). The exact mechanism of apocrine secretion of CLD is not well understood, but the process is known to incorporate portions of the apical plasma membrane, which includes proteins that form the CLD docking complex, as well as cytoplasmic elements including secretory vesicles, as CLDs are enveloped for secretion as MFGs (Kurosumi, Kobayashi et al. 1968, Wu, Howell et al. 2000, Honvo-Houéto, Henry et al. 2016).

Studies in mice indicate that formation of the CLD docking complex limits the amounts of cytoplasm included in MFGs and enhances the efficiency of lipid secretion (Oftedal 2012). Further functions of the CLD docking complex remain to be explored, and we anticipate that it may act as an intracellular scaffold and/or signaling hub to regulate overall milk production and secretion.

Milk fat secretion is critical to lactation success. Genetic disruption of the CLD synthetic machinery and docking complex components leads to poor offspring growth or starvation and death due to low milk consumption (Vorbach, Scriven et al. 2002, Ogg, Weldon et al. 2004, Russell, Schaack et al. 2011, Wang, Lv et al. 2012, Monks, Dzieciatkowska et al. 2016, Zhao, Ke et al. 2020, Jeong, Kadegowda et al. 2021). Models targeting the CLD synthetic machinery drive low milk fat production and secretion, decreasing milk caloric content (Smith, Cases et al. 2000, Beigneux, Vergnes et al. 2006, Russell, Schaack et al. 2011, Wang, Lv et al. 2012, Suburu, Shi et al. 2014), and models targeting the MFG secretion machinery drive the production of extremely large and unstable MFGs which potentially clog the ductal network, blocking overall milk secretion (Vorbach, Scriven et al. 2002, Ogg, Weldon et al. 2004, Monks, Dzieciatkowska et al. 2016, Jeong, Kadegowda et al. 2021). These mechanistic details have been worked out in dairy animals and/or model organisms by electron microscopy, immunohistochemistry & fluorescence and most recently, by elegant intravital imaging of glandular tissue (Mather, Masedunskas et al. 2019). Mammary tissue is difficult to obtain from humans for use with these methods, however, and our understanding of the regulation of human milk lipid production and secretion is therefore limited. As the milk lipid biosynthetic and secretory machinery are known to be retained on MFGs, we aimed to expand our understanding of human milk fat synthesis and secretion using a quantitative untargeted proteomic approach in MFGs. We also aimed to directly compare our findings from human samples to murine samples to identify how well conserved the MFG production machinery is conserved between species, and therefore, how representative experimental murine models are of human milk fat production. This is a particularly relevant question because the wide disparity between milk fat content in humans (3-4%) (Ballard and Morrow 2013) and mice (>20%) (Görs, Kucia et al. 2009) could indicate divergent mechanisms of milk fat secretion.

Previous efforts to define the human MFGM have identified the presence of MFG synthesis and docking complex protein homologs, including CIDEA, PLIN2, XDH/XOR and multiple BTN family members (Cavaletto, Giuffrida et al. 2002, Fortunato, Giuffrida et al. 2003, Liao, Alvarado et al. 2011, Spertino, Cipriani et al. 2012, Yang, Zheng et al. 2015, Lu, Wang et al. 2016, Yang, Cong et al. 2016, Juvarajah, Wan-Ibrahim et al. 2018, Zhang, Jiang et al. 2021), in addition to over 400 other proteins. These studies have considered all proteins associated with MFGs to be membrane-bound proteins. However, due to the apocrine mechanism of milk lipid secretion, MFGs also consist of variable amounts proteins from the cytoplasmic compartments (Patton and Huston 1988). Distinguishing these from true membrane-bound proteins can clarify which are required for CLD docking, envelopment and MFG secretion. We therefore directly compared the MFG proteome and the fractionated MFGM proteome from 13 women across early to midlactation. To isolate MFGMs, we utilized physical disruption rather than detergent-based disruption, as others have used for human MFGM analysis, to limit the solubilization and loss of individual proteins from membrane-bound complexes and maximize their isolation by centrifugation. Advancements in proteomics technology have allowed us to identify a far greater number of MFG and MFGM proteins than have previously been identified and our pathway analyses point toward potential regulators of milk fat synthesis and secretion.

## Materials and Methods

### Human and mouse milk collection

Human milk (<1oz) was collected after an overnight fast in postpartum women, as part of a randomized controlled trial of diet composition in the control of gestational diabetes (Clinical Trials #NCT02244814). Milk collection protocols were approved by the Colorado Multiple Institutional Review Board (protocol #14–1538) and all participants gave their informed consent, as previously described (Martin Carli, Trahan et al. 2020). Participants visits occurred at 2 weeks (5 samples), 2 months (3 samples) or 4-5 months postpartum (5 samples), following term deliveries (≥ 37 wks). Milk was collected from a total of 9 participants.

Two of these provided samples at both the 2 wk and 4-5 mo timepoints, and one participant provided milk at all three timepoints. Milk collections were not standardized with respect to the time of infant feeding or pumping. Samples were placed on ice in a cooler and transported to a study visit at the Clinical Translational Research Center at the University of Colorado Anschutz Medical Campus and then brought to the laboratory.

Mouse milk was collected from primiparous CD1 females from breeding colonies maintained in the AAALAC-Accredited (#00235) Center for Comparative Medicine at the University of Colorado Anschutz Medical Campus. The colony was housed under 14:10 hr light:dark cycle at a temperature of 72±2°F, humidity of 40±10% and fed standard chow (Teklad/Envigo 2920X) and hyperchlorinated (2-5 ppm) water. Females were mated with CD1 males and then housed individually prior to parturition. The day a litter was first seen was counted as lactation day 1 (L1). Dams were allowed to nurse their natural born litter (litters were not standardized, avg: 12±2 pups/litter). Milk samples were collected from 18 dams at L9-11, after 3 hrs of separation from pups, as previously described (Monks, Orlicky et al. 2022). Briefly, intraperitoneal (IP) xylazine was given at a dose of 8 mg/kg. When the mouse was relaxed enough to have ceased ambulation around the cage (about 5 min), the milking procedure was initiated. The mouse was picked up, and with gentle hand-restraint, a single dose of oxytocin (0.25 IU, 0.25 ml in sterile saline) was given IP. Milk let-down occurred within 1 min and milk removal was started. Our standard milking apparatus, attached to house vacuum, was used. Hand restraint was used throughout the milking procedure. Milk was collected and processed at room temperature to avoid changes in protein segregation between phases. All animal experiments and procedures were approved by the University of Colorado School of Medicine’s Institutional Animal Care and Use Committee on protocol 00985 (PI: McManaman).

### MFG and MFGM isolation

MFGs from both human and murine milk samples were isolated according to procedures previously described (Monks, Orlicky et al. 2022) which were informed by established methods (Patton and Huston 1986, Wu, Howell et al. 2000, Monks, Dzieciatkowska et al. 2016). Briefly, fresh whole milk was gently combined with ∼10 vol of PBS and centrifuged at 1,500 xg at room temperature. The mouse milk and two human samples were mixed 1:1 with 10% sucrose and layered under PBS for this first wash. The remaining human samples were not mixed with 10% sucrose, as this appeared to contribute to sample loss. Floated MFG were collected, washed twice by centrifugation through PBS and stored at −80°C. Membranes were isolated from intact human MFG using procedures described for isolation of bovine MFGM (Mather 2000). Briefly, frozen MFG were thawed on ice, Dounce homogenized (100 strokes) and centrifuged at 22,000 xg for 20 min at 4°C. Pelleted membranes (MFGM) were stored at −80°C prior to proteomic analysis.

### Liquid chromatography – tandem mass spectrometry (LC-MS/MS)

Isolated MFG or MFGM samples were precipitated with 10% trichloroacetic acid for 2 hr at −20°C. The precipitated protein samples were pelleted by centrifugation at 14,000 xg for 20 min at 4°C, rinsed in ice-cold acetone and centrifuged again. The pellet was air-dried and solubilized in 5% SDS, 100 mM DTT in 100 TEAB. The samples were digested using the S-Trap filter (Protifi, Huntington, NY) according to the manufacturer’s procedure. Briefly, samples were reduced with 10 mM DTT at 55°C for 30 min, cooled to room temperature, and then alkylated with 25 mM iodoacetamide in the dark for 30 min. Next, a final concentration of 1.2% phosphoric acid and then six volumes of binding buffer (90% methanol; 100 mM triethylammonium bicarbonate, TEAB; pH 7.1) were added to each sample. After gently mixing, the protein solution was loaded to a S-Trap filter, spun at 1,000 xg for 1 min, and the flow-through collected and reloaded onto the filter. This step was repeated three times, and then the filter was washed with 200 μL of binding buffer 3 times. Finally, 1 μg of sequencing-grade trypsin (Promega) and 150 μL of digestion buffer (50 mM TEAB) were added onto the filter and digestion was carried out at 37°C for 6 hr. To elute peptides, three stepwise buffers were applied, with 100 μL of each with one more repeat, including 50 mM TEAB, 0.2% formic acid in H_2_O, and 50% acetonitrile and 0.2% formic acid in H2O. The peptide solutions were pooled, lyophilized and resuspended in 100 μl of 0.1% FA.

20 μl of each sample was loaded onto individual Evotips for desalting and then washed with 20 μL 0.1% FA followed by the addition of 100 μL storage solvent (0.1% FA) to keep the Evotips wet until analysis. The Evosep One system (Evosep, Odense, Denmark) was used to separate peptides on a Pepsep column, (150 um inter diameter, 15 cm) packed with ReproSil C18 1.9 um, 120A resin. The system was coupled to the timsTOF Pro mass spectrometer (Bruker Daltonics, Bremen, Germany) via the nano-electrospray ion source (Captive Spray, Bruker Daltonics). The mass spectrometer was operated in PASEF mode (TIMS/PASEF). The ramp time was set to 100 ms and 10 PASEF MS/MS scans per topN acquisition cycle were acquired. MS and MS/MS spectra were recorded from m/z 100 to 1700. The ion mobility was scanned from 0.7 to 1.50 Vs/cm^2^. Precursors for data-dependent acquisition were isolated within ±1 Th and fragmented with an ion mobility-dependent collision energy, which was linearly increased from 20 to 59 eV in positive mode. Low-abundance precursor ions with an intensity above a threshold of 500 counts but below a target value of 20,000 counts were repeatedly scheduled and otherwise dynamically excluded for 0.4 min. TIMS/PASEF mass spectrometry has been shown to be an ultra-sensitive proteomics approach (Meier, Park et al. 2021).

### Database searching and protein identification

MS/MS spectra were extracted from raw data files and converted into .mgf files using MS Convert (ProteoWizard, Ver. 3.0). Peptide spectral matching was performed with Mascot (Ver. 2.6) against the Uniprot h database. Mass tolerances were ±15 ppm for parent ions, and ±0.4 Da for fragment ions. Trypsin specificity was used, allowing for 1 missed cleavage. Met oxidation, protein N-terminal acetylation and peptide N-terminal pyroglutamic acid formation were set as variable modifications with Cys carbamidomethylation set as a fixed modification.

Scaffold (version 5.0, Proteome Software, Portland, OR, USA) was used to validate MS/MS based peptide and protein identifications. Peptide identifications were accepted if they could be established at greater than 95.0% probability as specified by the Peptide Prophet algorithm.

Protein identifications were accepted if they could be established at greater than 99.0% probability and contained at least two identified unique peptides.

### Statistical analyses

Milk collections were treated as independent samples, even though a subset of participants provided repeated samples. This is because milk fat composition is largely affected by diet and collection variables that we could not account for, such as whether foremilk or hindmilk was collected, or time since last breast emptying. Statistical analyses were conducted as described using Graphpad Prism 9.5.1 and Metaboanalyst 5.0 (https://www.metaboanalyst.ca/home.xhtml).

### Pathway analyses

Gene symbols of the proteins detected in ≥ 50% of MFG or MFGM samples, for human, or ≥ 50% of murine MFG samples were utilized for pathway enrichment analysis using Metascape (https://metascape.org/gp/index.html#/main/step1). This tool queries multiple different ontology sources, and minimizes redundancy by clustering related pathway terms (Zhou, Zhou et al. 2019). We first identified all statistically enriched terms (can be GO/KEGG terms, canonical pathways, hall mark gene sets, etc., based on the default choices under Express Analysis), accumulative hypergeometric p-values and enrichment factors were calculated and used for filtering. Remaining significant terms were then hierarchically clustered into a tree based on Kappa-statistical similarities among their gene memberships (similar to what is used in NCI DAVID site). Then 0.3 kappa score was applied as the threshold to cast the tree into term clusters. To visualize our results in the context of cellular mechanisms, we utilized Reactome’s (Fabregat, Sidiropoulos et al. 2018) pathway diagram viewer (https://reactome.org/, version 3.7) with our gene symbol lists.

## Results

In this study, we defined quantitative proteomic profiles of MFG and MFGM pairs from 13 women across early (2 weeks postpartum, n=5) to mature lactation (2 months postpartum, n=3, and 4-5 months postpartum, n=5) by LC-MS/MS. The sum of the total spectral counts was not different between MFG and MFGM proteins, indicating effective sample preparation and similar loading between sample types (**Fig. 1a**). We included in our analyses only the 2,257 proteins which were detected in ≥ 50% of replicates in one or both groups. We considered the possibility that MFG and/or MFGM proteins from late-transitional/early mature milk at 2 wk postpartum could differ from those found in mature milk from 2-5 mo postpartum, especially as bovine MFGM-bound proteins have been shown to change from colostrum to mature milk (Reinhardt and Lippolis 2008). By principal components analysis (PCA), we found that MFG proteins (**Fig. 1b**) could not be distinguished across timepoints by the first five principal components (80.6% of total variance). MFGM proteins at transitional vs mature timepoints were separated across PC2 (13.8%; **Fig. 1c**). BTN1A1 was the biggest driver of this separation, with a loading score of 0.23 (**Fig. 1d**). The abundance of BTN1A1 increased from 0.027±0.002 normalized spectral abundance factor (NSAF) in early lactation to 0.035±0.004 NSAF in mature lactation (**Fig. 1e**; mean ±SD, adj. p<0.01). Other components of the CLD docking complex, XDH/XOR, PLIN2 and CIDEA, which are increased in the transition from bovine colostrum to mature milk (Reinhardt and Lippolis 2008), were not increased during the transition from early to mature human lactation (**Fig. 1e**). When the entire proteomic dataset was considered, we did not find statistical differences between early and mature lactation (**Sup. Fig. 1**). Many factors, including time since last feeding and time respective to the feeding/pumping bout are expected to affect CLD docking and envelopment (Masedunskas, Chen et al. 2017, Mather, Masedunskas et al. 2019) and therefore, protein composition of the MFGM. We are unable to account for these collection details in the current study, which did not standardize these factors, and we do not have access to samples from the colostrum phase. As we were underpowered to investigate temporal changes across lactation, we combined all timepoints for subsequent analyses to maximize our statistical power.

**Figure 1.**
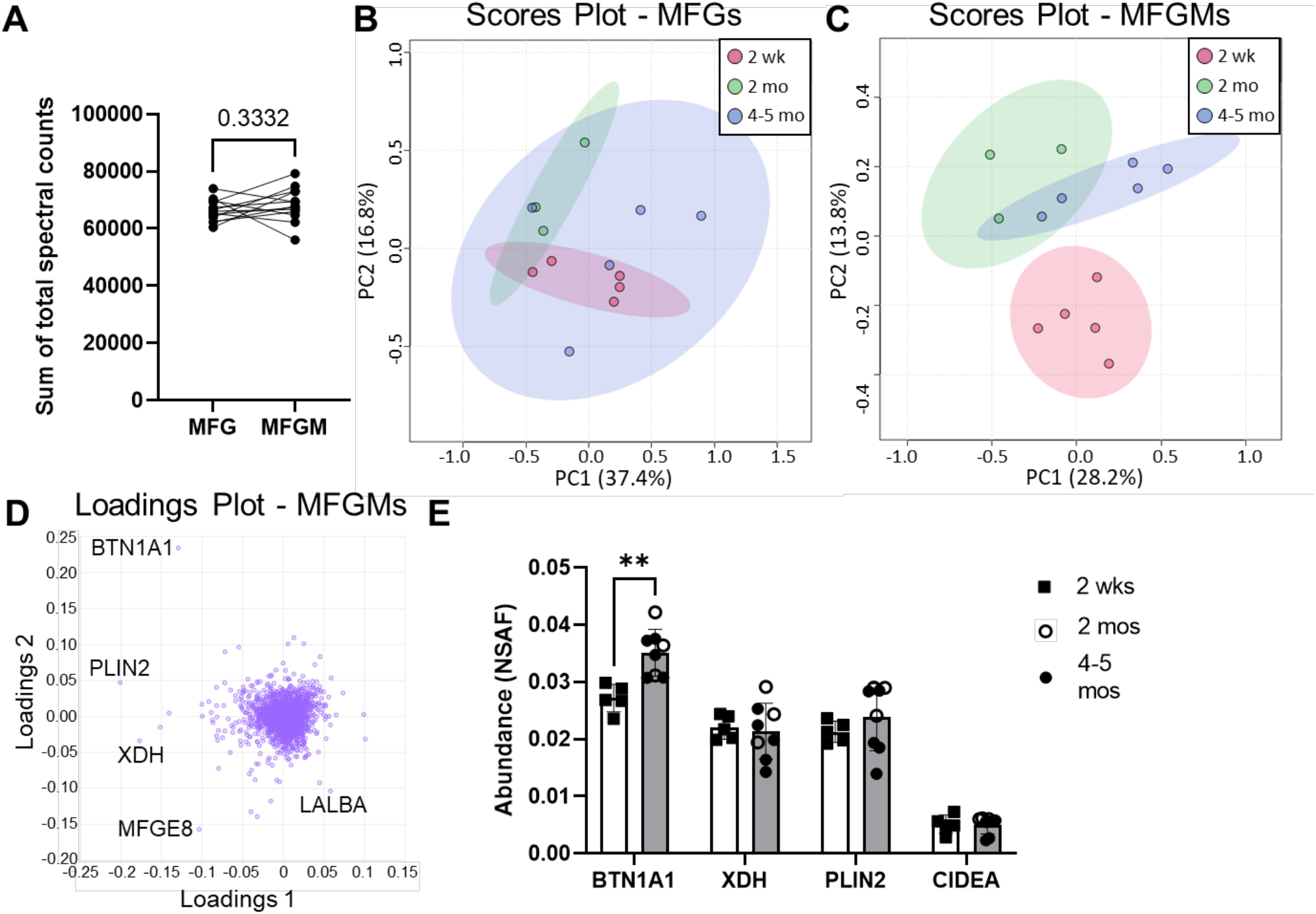
Human MFG and MFGM protein abundance over time. Sum of human MFG and MFGM spectral counts by LC-MS/MS (**A**). Principal components analysis of 1,824 MFG-associated (**B**) and 2,120 MFGM-associated (**C**) proteins separated by time postpartum: red=2wk, green=2mo, blue=4-5mo. Pareto scaling was used to limit the effects of large fold changes found in low abundance proteins. Loadings of MFGM-associated protein PCA (**D**). Abundance of known CLD docking proteins BTN1A1, XDH, PLIN2 and CIDEA (**E**). Open bars=2wks, gray bars=2-5mo, open circles=2mo, closed circles=4mo. Multiple nonparametric (Mann-Whitney), unpaired t-tests, adjusted for multiple testing with the Holm-Šídák method. **p<0.01.

We identified 1,687 proteins that were common between the two sample types, as well as 137 unique proteins in the MFG samples and 433 unique proteins in the MFGM samples. (**Fig. 2a**). Stoichiometric analysis of proteomic data is challenging as sequence coverage is inconsistent due to differences in tryptic sites and ionization efficiency. However, we obtained high coverage (>60%) of proteins implicated in CLD docking (BTN1A1, XDH/XOR, PLIN2 (Monks, Orlicky et al. 2022) (**Sup. Fig. 2a**). We find that of CLD docking proteins, both in MFG samples and isolated MFGM proteins, BTN1A1 is the most abundant protein, followed by PLIN2 and XDH/XOR. These three proteins are more highly abundant than CIDEA, which is also implicated in CLD docking (Monks, Dzieciatkowska et al. 2016). A heatmap of the 20 most abundant proteins across both sample types is shown in **Fig. 2b**. We next compared MFG and MFGM proteins by PCA and identified distinctly different proteomes (**Fig. 2c**). As expected, the loadings contributing to this distinction primarily consisted of proteins implicated in CLD docking; XDH/XOR, BTN1A1, PLIN2 and CIDEA (**Fig. 2d**). To identify possible additional components of the human CLD docking complex, we identified differential protein abundance between sample types, or MFGM enrichment, by volcano plots, comparing both differences (**Fig. 2e**) to assess highly abundant proteins, and fold changes (**Fig. 2f**) to assess proteins with low abundance.

**Figure 2.**
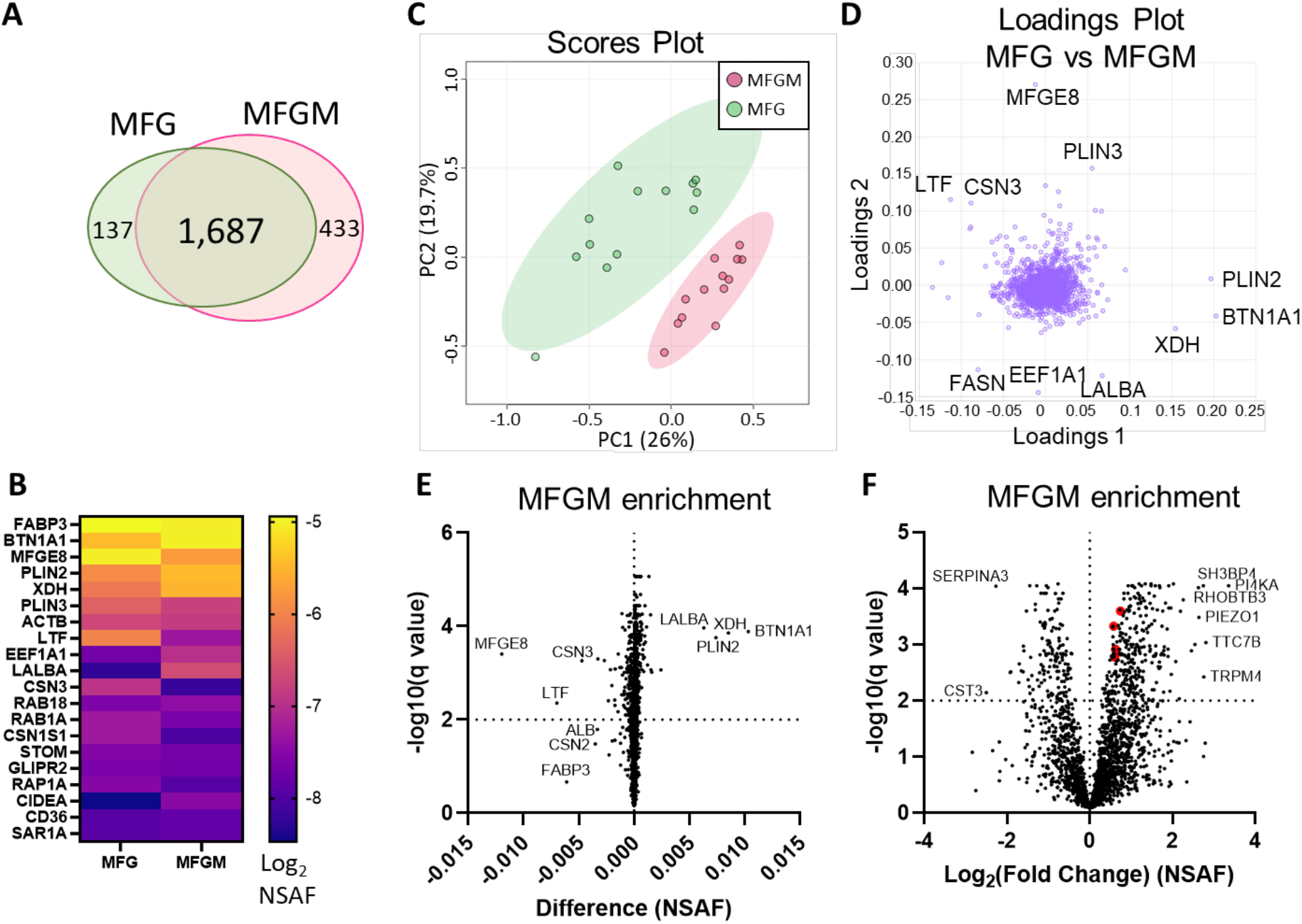
Differential protein abundance between human MFGs and MFGMs. Venn diagram of shared and distinct human MFG- and MFGM-associated proteins (**A**). Abundance of the top 20 human MFG- and MFGM-associated proteins (**B**, Log_2_ NSAF), sorted by median of MFG and MFGM values. Principal components analysis of MFG-associated (green) and MFGM-associated (red) proteins (**C**). Pareto scaling was used to limit the effects of large fold changes found in low abundance proteins. Loadings of MFG- and MFGM-associated protein PCA (**D**). Multiple paired t-tests of MFG-vs MFGM-associated proteins, calculated by difference (MFGM-MFG, **E**) or by Log_2_ fold change (**F**) with FDR correction for multiple comparisons (1%). BTN1A1, XDH, PLIN2 and CIDEA are labeled red in F.

We conducted pathway analysis using Metascape, which comprehensively utilizes KEGG Pathway, GO Biological Processes, Reactome Gene Sets, Canonical Pathways, CORUM, WikiPathways, and PANTHER Pathway as ontology sources. Differentially abundant proteins identified in **Fig. 2e** were separated into MFGM-enriched and MFG-enriched protein lists.

Pathway enrichment analysis of MFGM-enriched proteins (n=628; **Fig. 3a, Sup. Fig. 3**) identified pathway clusters related to lipid metabolism (lipid biosynthetic process and regulation of lipid metabolic process), endoplasmic reticulum (ER; response to ER stress, protein processing in the ER, protein N-linked glycosylation) and intracellular trafficking, as expected. Intracellular trafficking-related pathway clusters include transport of small molecules, membrane trafficking, intracellular protein transport, lipid localization, localization within membrane, endocytosis, regulation of secretion, regulation of vesicle-mediated transport and export from cell. Interestingly, this analysis also identified neutrophil degranulation and VEGFA-VEGFR2 signaling pathway clusters.

**Figure 3.**
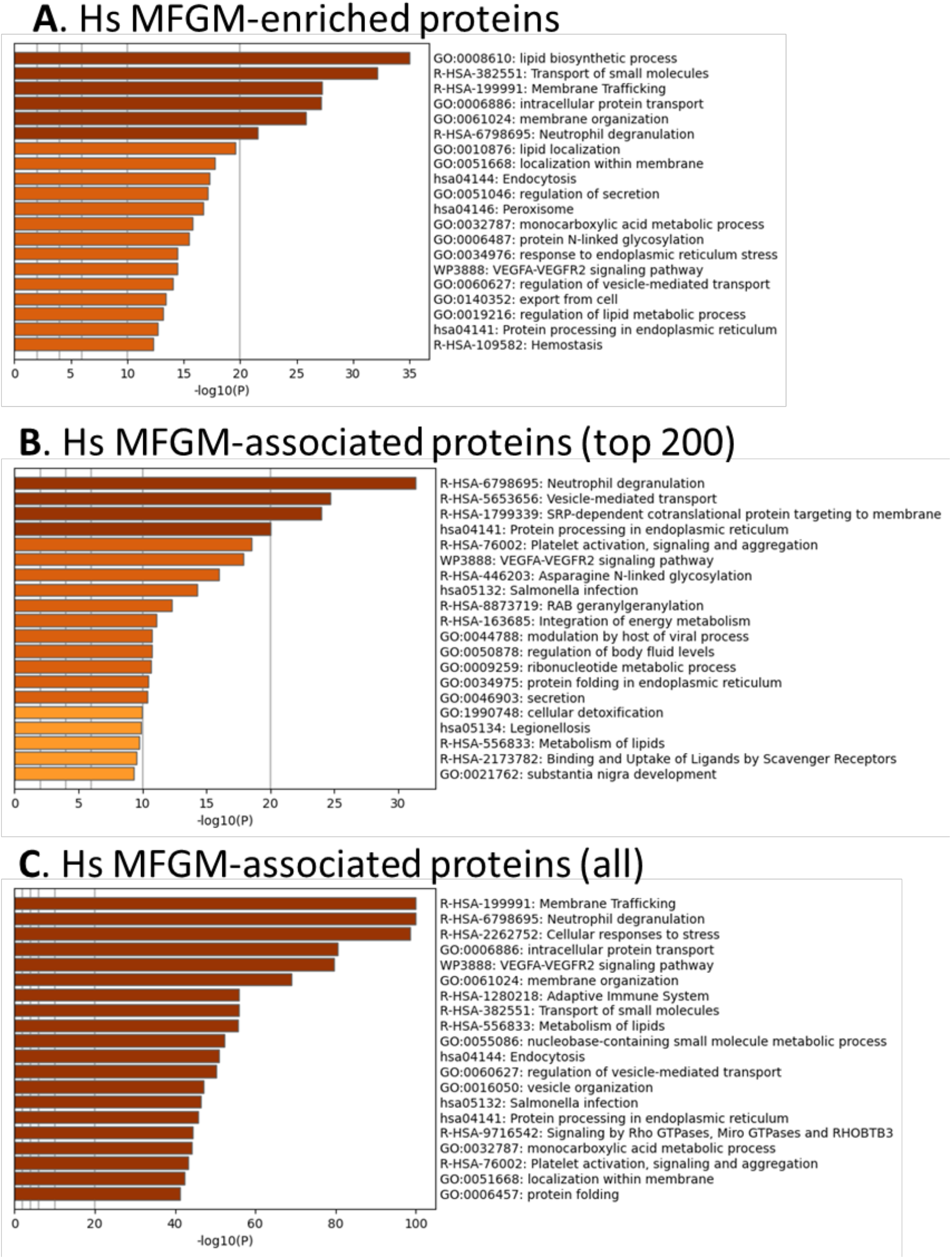
Metascape pathway analyses of human MFGM-enriched and MFGM-associated proteins. Pathway enrichment analysis of MFGM-enriched proteins, calculated by difference (**A**). Pathway enrichment analysis of the top 200 (sorted by median) MFGM-associated proteins (**B**). Pathway enrichment analysis of all MFGM-associated proteins (**C**).

Not all MFGM-associated proteins were statistically significantly different in comparison to the whole MFG, due to similar distribution across both membrane and cytoplasmic compartments. We considered the possibility that any MFGM-associated protein, both the highly abundant and less abundant proteins, could provide insight into the cellular processes regulating MFG synthesis and secretion. We therefore conducted pathway analyses with the most highly abundant (n=top 200; **Fig. 3b, Sup. Fig. 4**) and all (n=2,120; **Fig. 3c, Sup. Fig.5**) MFGM-associated proteins, without comparison to MFG-associated proteins. Lipid, ER and transport-related clusters as well as neutrophil degranulation and VEGFA-VEGFR2 signaling pathway clusters were identified in both analyses. RAB geranylgeranylation and signaling by Rho GTPases, Miro GTPases and RHOBTB3 clusters were also identified by these analyses.

**Figure 4.**
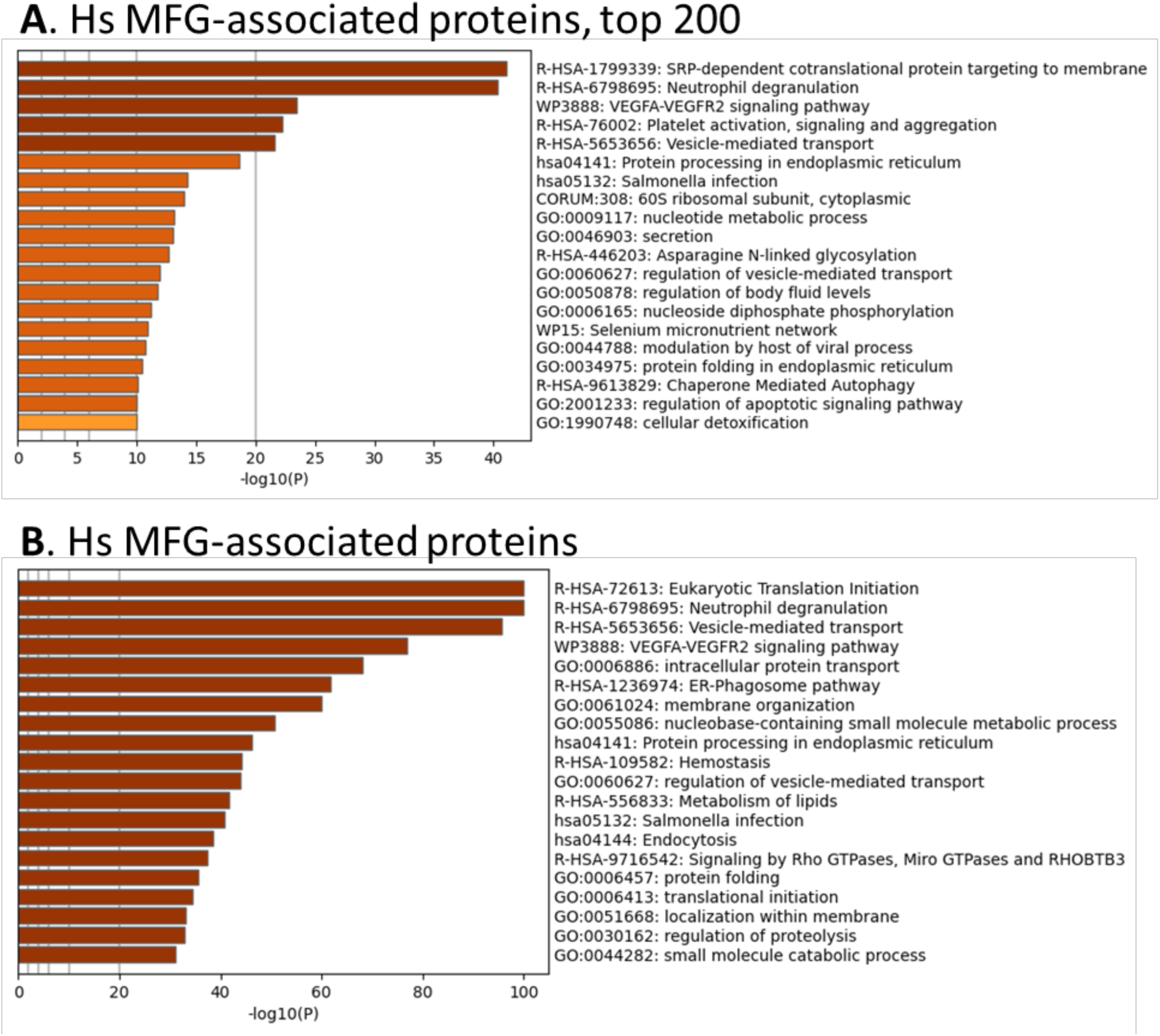
Metascape pathway analyses of human MFG-associated proteins. Pathway enrichment analysis of the top 200 (sorted by median) MFG-associated proteins (**A**). Pathway enrichment analysis of all MFG-associated proteins (**B**).

Although the MFG and MFGM proteomes were distinct by principal components analysis, we aimed to determine which pathways were found in common between MFG-associated and MFGM-associated proteins and which were distinct. Using both the most abundant (**Fig. 4a, Sup. Fig. 6**) and all (**Fig. 4b, Sup. Fig.7**) MFG-associated proteins for pathway analysis, we identified ribosome-related pathway clusters. These include eukaryotic translation initiation, SRP-dependent cotranslational protein targeting of membrane and 60S ribosomal subunit, cytoplasmic. This is consistent with the inclusion of cytoplasmic fragments in MFGs that have not been further purified to yield MFGM isolates. However, we also identified pathway clusters which we identified using the MFGM proteome, including neutrophil degranulation, VEGFA-VEGFR2 signaling, vesicle-mediated transport and Rho GTPases, Miro GTPases and RHOBTB3, in addition to lipid metabolism, endoplasmic reticulum and intracellular trafficking-related pathway clusters. Therefore, pathway analysis of MFG-associated pathways identifies pathways active both in the endoplasmic reticulum and apical plasma membranes as well as the neighboring cytoplasm.

As we and others utilize murine models to investigate mechanisms supporting milk secretion by the mammary gland, we aimed to identify the similarities between mouse and human MFG secretion. A fuller understanding of how well the mouse mammary gland represents the human breast can inform translational research into human lactation. We therefore collected milk from lactating CD1 mouse dams and isolated MFGs. The small volumes available precluded us from isolating the MFGM fraction from these samples, and therefore, we were restricted to analyzing the MFG proteome, with the understanding that these proteins comprise a mix of cytoplasmic and membrane-bound proteins. We obtained similar coverage of the known CLD docking components (>50-60%; **Sup. Fig. 2b**). Interestingly, in the murine samples, Xdh/Xor was more highly abundant than Btn1a1 and Cidea, and these three were more highly abundant than Plin2 (**Fig. 5a**). Pathway analyses of murine MFG-associated proteins identified similar pathway clusters as the human analysis, with ribosome, endoplasmic reticulum, lipid metabolism and intracellular transport pathway clusters being well represented (**Fig. 5b**). Of the notable pathways identified in the human analysis described above, we also identified neutrophil degranulation, Rho GTPases, Miro GTPases and RHOBTB3, and RAB geranylgeranylation with murine MFG-associated proteins (**Sup. Fig 8**). Notably missing from the murine pathway cluster list was VEGFA-VEGFR2 signaling, which was highly significant in the human pathway analyses.

**Figure 5.**
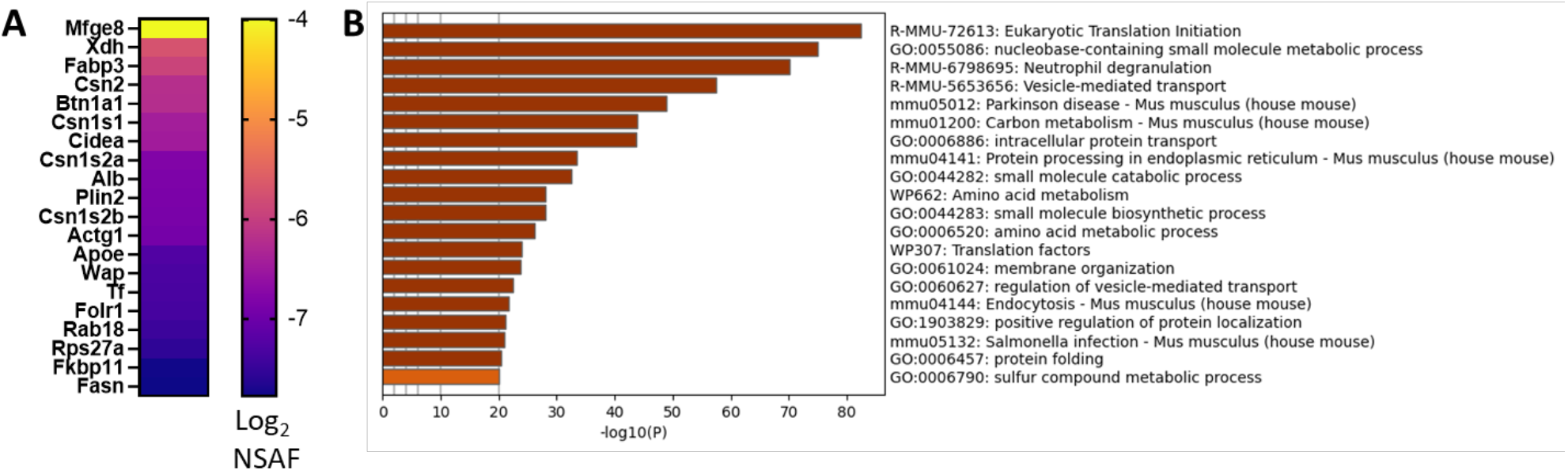
CD1 mouse MFG proteome at L10. Abundance of the top 20 MFG-associated proteins (**A**, Log_2_ NSAF), sorted by median. Metascape pathway enrichment analyses of MFG-associated proteins (**B**).

**Figure 6.**
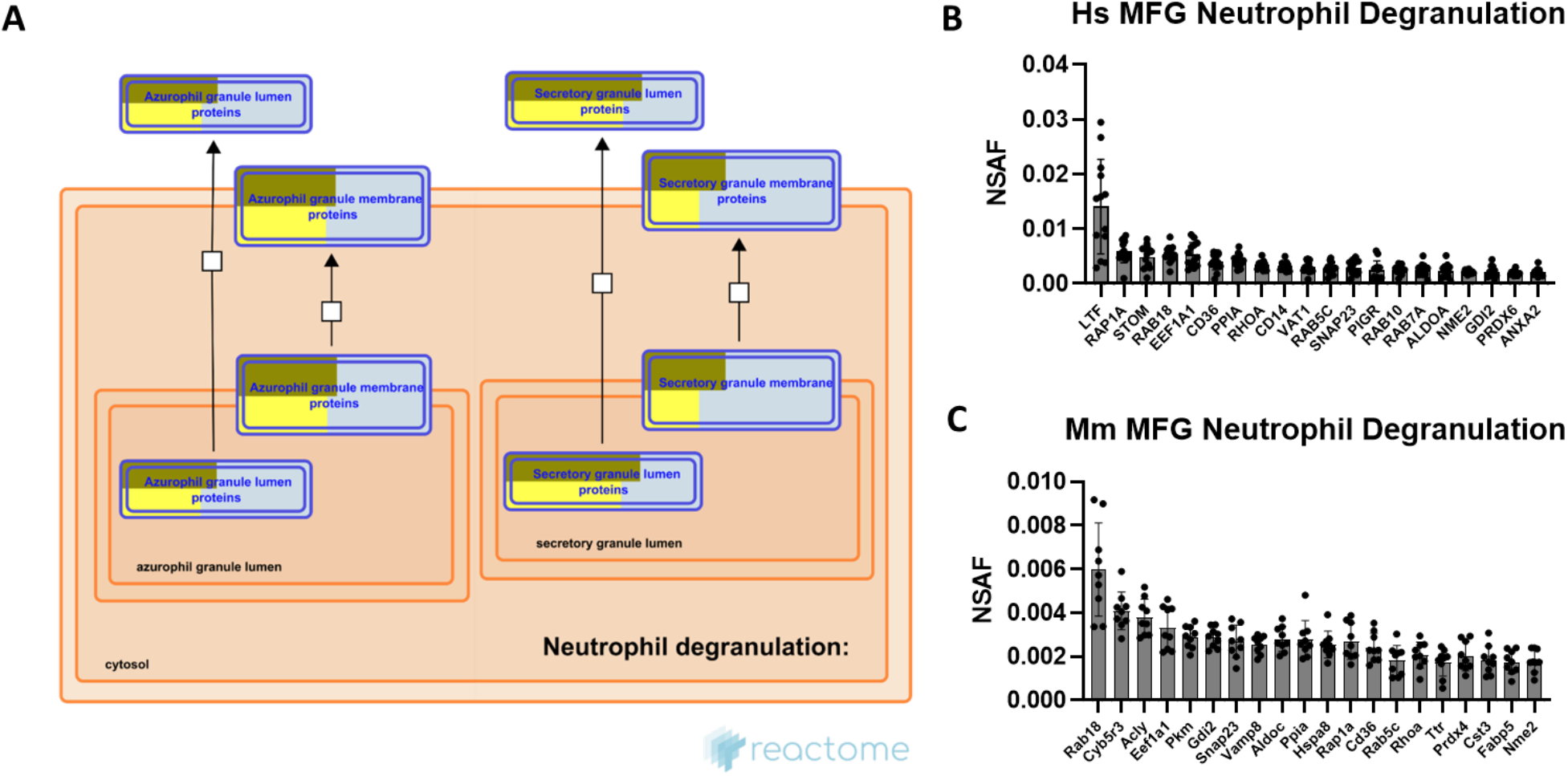
Pathway enrichment analysis: Neutrophil degranulation. Reactome pathway diagram with overlaid overrepresentation pathway enrichment analyses of human and mouse MFG-associated proteins in the neutrophil degranulation pathway, including only the primary (azurophil) and secretory granule entities (**A**). Entities are re-colored (gold for human, yellow for mouse) if they were represented in the submitted data set. The top 20 most highly abundant neutrophil degranulation-related proteins from the human (**B**) and mouse (**C**) MFG proteomes, sorted by median.

**Figure 7.**
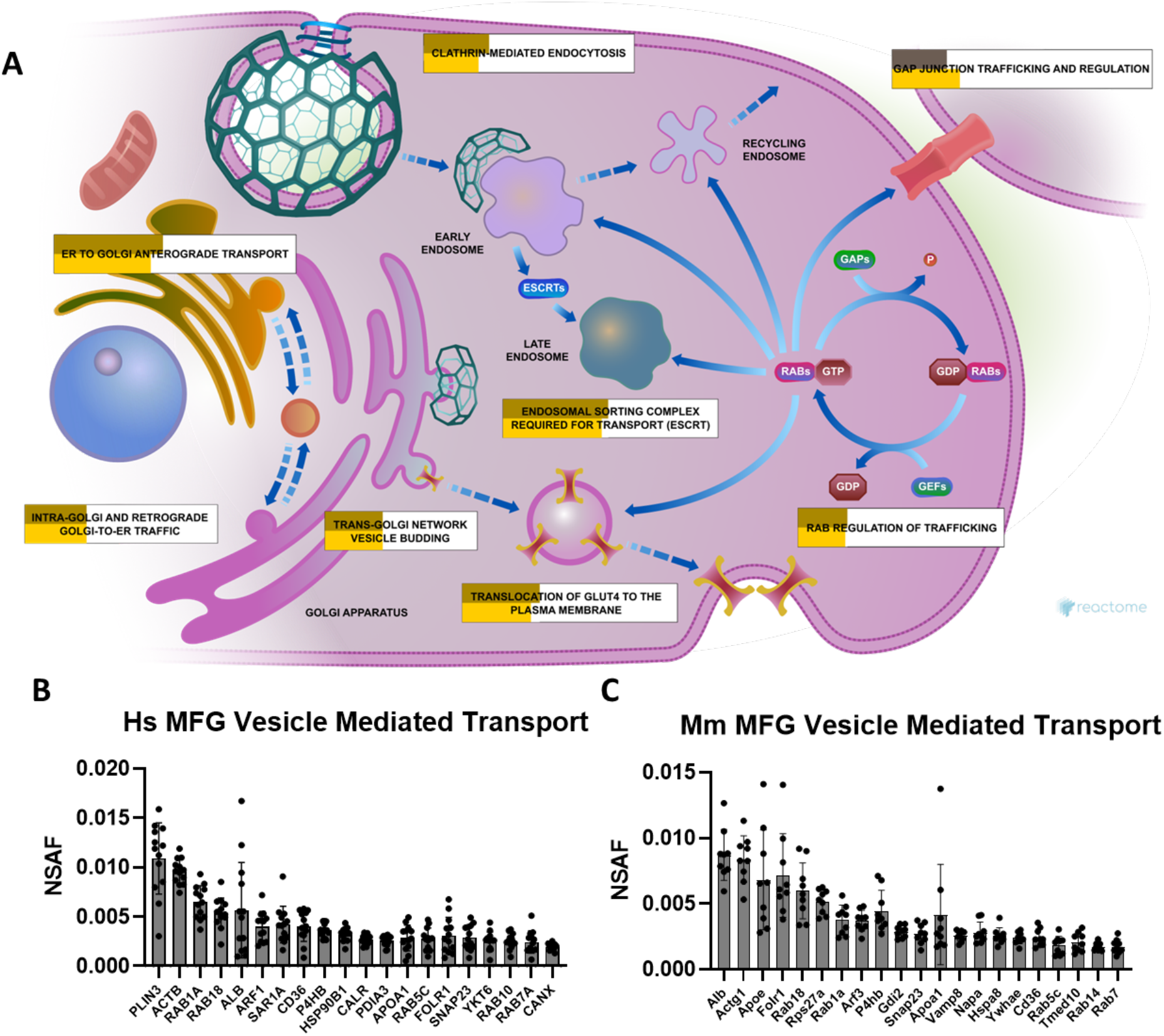
Pathway enrichment analysis: Vesicle mediated transport and membrane trafficking. Reactome pathway diagram with overlaid overrepresentation pathway enrichment analyses of human and mouse MFG-associated proteins in the membrane trafficking pathway (**A**). Entities are re-colored (gold for human – upper, yellow for mouse – lower, gray if not statistically significant) if they were represented in the submitted data set. The top 20 most highly abundant vesicle mediated transport-related proteins from the human (**B**) and mouse (**C**) MFG proteomes, sorted by median.

**Figure 8.**
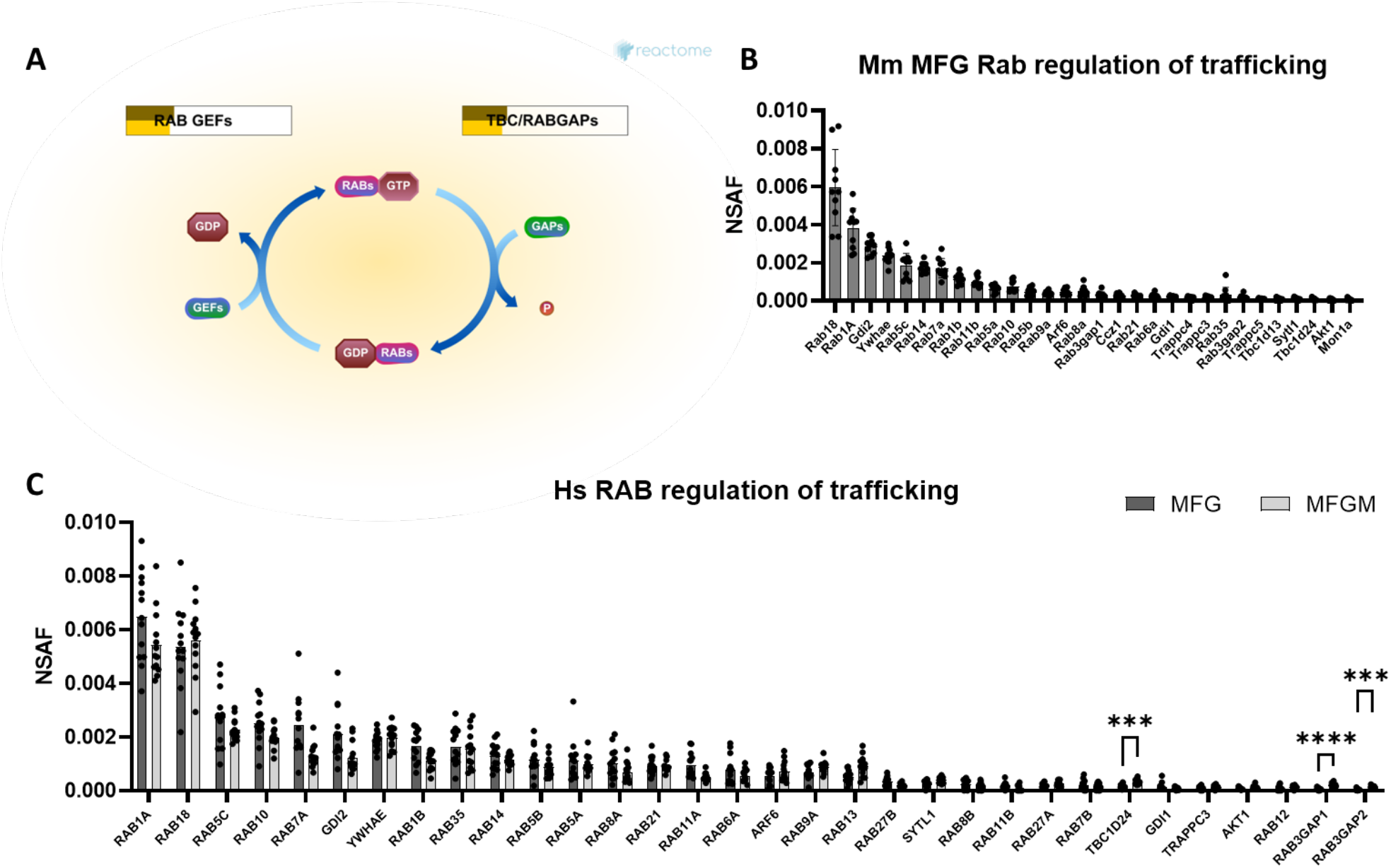
Pathway enrichment analysis: RAB regulation of trafficking. Reactome pathway diagram with overlaid overrepresentation pathway enrichment analyses of human and mouse MFG-associated proteins in the RAB regulation of trafficking pathway (**A**). Entities are re-colored (gold for human – upper, yellow for mouse – lower) if they were represented in the submitted data set. All RAB regulation of trafficking-related proteins found in the mouse MFG (**B**) and human MFG (dark gray) and MFGM (light gray, **C**) proteomes, both sorted by MFG median.

Of the pathways found in common between human and mouse MFG-associated protein pathway analyses, neutrophil degranulation was consistently one of the most highly significant. As it was only recently discovered that MFG secretion is stimulated upon oxytocin-mediated myoepithelial cell contraction (Masedunskas, Chen et al. 2017), the processes regulating this stimulated form of secretion remain largely unknown. Therefore, commonalities with the carefully controlled neutrophil degranulation process may provide insight into stimulated MFG secretion mechanisms. These two secretory processes are structurally distinct, with neutrophil degranulation constituting exocytosis of secretory vesicles, whereas MFG secretion involves the envelopment of the CLD in apical plasma membrane and potentially other vesicle membranes. Neutrophils contain at least 4 distinct types of preformed secretory vesicles known as granules, which are thought to form sequentially during neutrophil differentiation, and which contain distinct effector subsets (Le Cabec, Cowland et al. 1996). These include primary, secondary, tertiary and secretory granules, with secondary granules containing lactoferrin, an antibacterial protein which is also highly abundant in milk (Kell, Heyden et al. 2020). Pathway analysis of both human (**Sup. Fig. 9a**) and mouse (**Sup. Fig. 9b**) MFG-associated proteins identified the strongest overrepresentation of primary (azurophilic) granule membrane proteins and secretory granule lumen proteins, as illustrated by Reactome (**Fig. 6a**).

**Figure 9.**
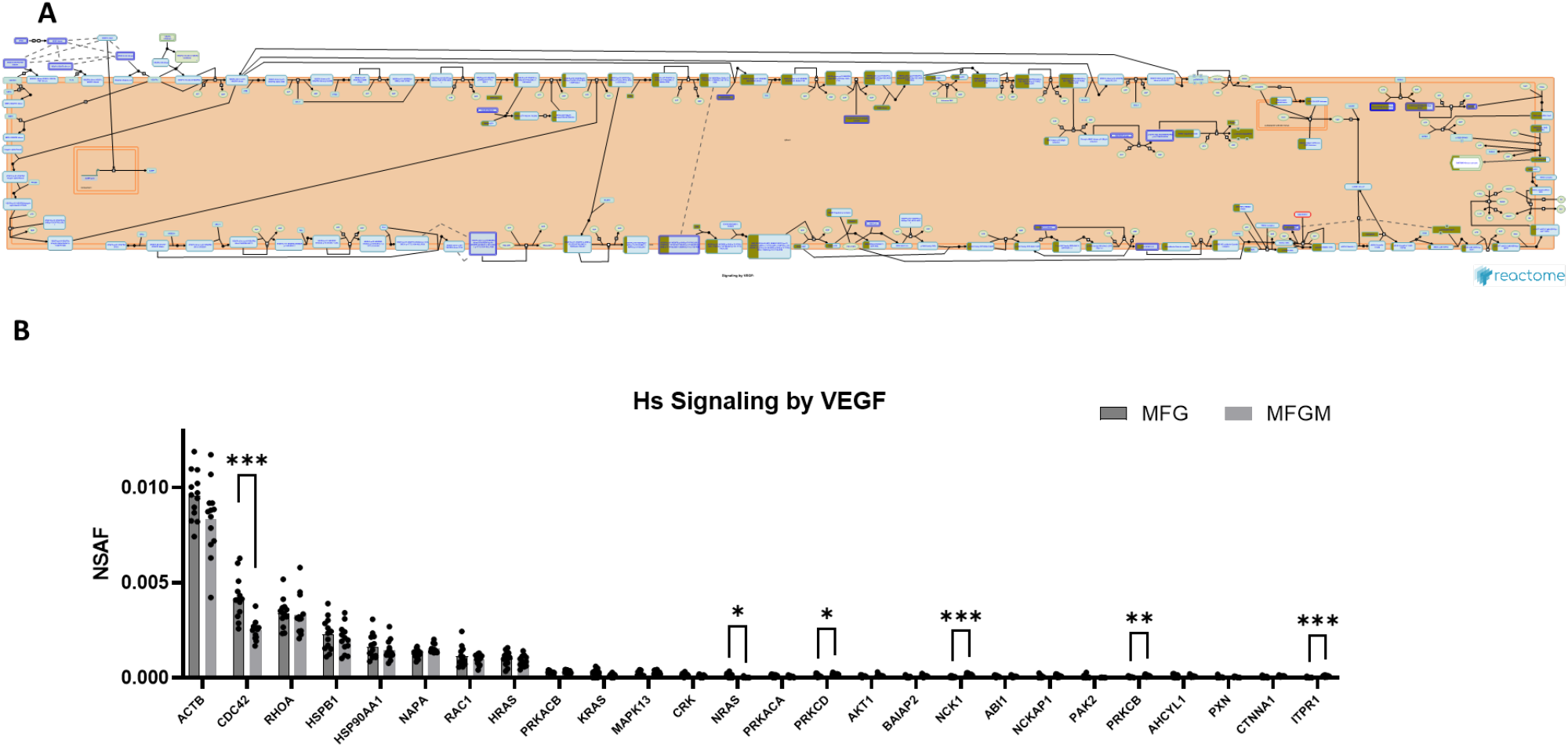
Pathway enrichment analysis: Signaling by VEGF. Reactome pathway diagram with overlaid overrepresentation pathway enrichment analyses of human MFG-associated proteins in the signaling by VEGF pathway (**A**). Entities are re-colored (gold) if they were represented in the submitted data set. All signaling by VEGF-related proteins found in the human MFG (dark gray) and MFGM (light gray) proteomes (**B**), both sorted by MFG median.

Neutrophil degranulation entails cytoskeletal remodeling to allow for granule trafficking to the plasma membrane, granule tethering and docking, granule priming for fusion, and fusion of the granule with the plasma membrane to allow for release of its contents. Small GTPases, which are critical for cytoskeletal remodeling and vesicle trafficking, are known to regulate many of the interactions regulating these processes (Lacy 2006). The top 20 most abundant neutrophil degranulation pathway proteins identified in human and mouse MFG-associated proteomes are shown in **Fig. 6b&c**, respectively. Common between these are the small GTPases RAP1A, member of RAS oncogene family (RAP1A), RAB18, member RAS oncogene family (RAB18), which is known in other cell types to regulate lipid droplet growth (Ozeki, Cheng et al. 2005, Xu, Li et al. 2018, Deng, Zhou et al. 2021), ras homolog family member A (RHOA), RAB5C, member RAS oncogene family (RAB5C), as well as the GDP dissociation inhibitor 2 (GDI2).

Also highly abundant is synaptosome associated protein 23 (SNAP23), which has been shown to form a soluble N-ethylmaleimide sensitive factor attachment protein receptor (SNARE) complex with vesicle-associated membrane protein 8 (Vamp8) in murine mammary epithelial cells, potentially allowing secretory vesicles to contribute membrane to the MFGM in addition to the apical plasma membrane (Honvo-Houéto, Henry et al. 2016).

Related to the specialized trafficking responsible for neutrophil degranulation, we also identified vesicle mediated transport, and its subcluster membrane trafficking, as overrepresented pathway clusters in both human (**Sup. Fig. 10**) and mouse (**Sup. Fig. 11**) MFG-associated proteomes. As illustrated by Reactome, multiple pathways were represented by the membrane trafficking pathway cluster which were significantly overrepresented in our MFG datasets, including ER to Golgi anterograde transport, intra-Golgi and retrograde Golgi-to-ER traffic, trans-Golgi network vesicle budding, translocation of GLUT4 to the plasma membrane, RAB regulation of trafficking, clathrin mediated endocytosis and endosomal sorting complex required for transport (ESCRT). Gap junction trafficking and regulation was also significantly overrepresented, but only in our murine dataset. The top 20 most abundant vesicle mediated pathway proteins identified in human MFG and MFGM- and mouse MFG-associated proteomes are shown in **Fig. 7b&c**, respectively. These again include RAB18, RAB5C and SNAP23, in addition to RAB1A, member RAS oncogene family (RAB1A), RAB10, member RAS oncogene family (RAB10) and RAB7A, member RAS oncogene family (RAB7A). Also common between species are actin beta (ACTB) and albumin (ALB).

The RAB family of small GTPases regulates many steps of intracellular membrane trafficking events, including vesicle formation, vesicle fusion with membranes, and vesicle movement along cytoskeletal networks. RAB proteins are peripheral membrane proteins, anchored to a membrane via a covalently attached lipid moiety (geranylgeranyl-, prenyl-, acetyl-group), and are regulated by effector proteins including escort, guanine nucleotide exchange factor (GEF), and GTPase-activating proteins (GAP)(Liu, Bartz et al. 2007, Kjos, Vestre et al. 2018, Homma, Hiragi et al. 2021). RAB18, in particular, has been shown to be involved in lipid droplet interactions with other organelles(Li, Luo et al. 2017, Zappa, Venditti et al. 2017). We anticipate that the collection of MFG RAB GEFs and RAB GAPs, overrepresented in analyses using Reactome (**Fig. 8a**), will provide insights into the cellular processes of MFG secretion. Most RAB proteins identified were found in both murine (**Fig. 8b**) MFGs and human (**Fig. 8c**) MFGs and MFGMs. These include RAB1A, RAB18, RAB5C, RAB10, RAB7A, GDI2, tyrosine 3-monooxygenase/tryptophan 5-monooxygenase activation protein epsilon (YWHAE), RAB1B, member RAS oncogene family (RAB1B), RAB35, member RAS oncogene family (RAB35), RAB14, member RAS oncogene family (RAB14), RAB5B, member RAS oncogene family (RAB5B), RAB5A, member RAS oncogene family (RAB5A), RAB8A, member RAS oncogene family (RAB8A), RAB21, member RAS oncogene family (RAB21), RAB6A, member RAS oncogene family (RAB6A), RAB9A, member RAS oncogene family (RAB9A), RAB11B, member RAS oncogene family (RAB11B), GDP dissociation inhibitor 1 (GDI1), RAB3 GTPase activating protein catalytic subunit 1 (RAB3GAP1) and RAB3 GTPase activating protein catalytic subunit 2 (RAB3GAP2). Also common to both species in the RAB regulation of trafficking pathway were ADP ribosylation factor 6 (ARF6), synaptotagmin like 1 (SYTL1), trafficking protein particle complex subunit 3 (TRAPPC3) and AKT serine/threonine kinase 1 (AKT1). Many RAB proteins have been implicated in progression of mammary carcinoma however, only RAB6A has been shown to play a role in lactation (Cayre, Faraldo et al. 2020).

Intriguingly, the VEGFA-VEGFR2 signaling pathway cluster was identified as highly significant in Metascape pathway analyses of the human MFG proteome (**Fig. 4**), but not the mouse MFG proteome (**Fig. 5**). We illustrate this pathway using Reactome in **Fig. 9a**, with the proteins identified in the human MFG and MFGM displayed in **Fig. 9b**. As vascular endothelial growth factor (VEGF) is best studied for its effects on endothelial cells, potential direct effects on epithelial cells are unexpected. We did not identify any VEGF receptors in either the human MFG or MFGM proteomes, indicating that if VEGF signaling is occurring in the mammary epithelial cells which secrete MFGs, it is not occurring on the apical surface, and is likely occurring across the basal membrane. Non-catalytic region of tyrosine kinase adaptor protein 1 (NCK1) and subunits of Protein Kinase A and C protein kinase cAMP-activated catalytic subunit beta (PRKACB), protein kinase cAMP-activated catalytic subunit alpha (PRKACA), protein kinase C delta (PRKCD) and protein kinase C beta (PRKCB) appear to be the main factors driving the identification of the VEGF signaling pathway. These signal transduction molecules are not specific to the VEGF signaling pathway, and we expect that their function in other signaling pathways active in the lactating mammary epithelial cells may be more relevant for understanding MFG synthesis and secretion.

## Discussion

Milk fat globules are unique, nutritionally important, membrane enveloped structures that are the primary source of neonatal calories and fat-soluble vitamins, fatty acids and lipid signaling molecules implicated in neonatal development (Oftedal 1984). We used ultra-sensitive TIMS/PASEF proteomics to quantify relative abundances of proteins in freshly isolated human MFG and their corresponding MFGM fractions and to compare mouse and human MFG proteomic profiles. Ours is the first study to comprehensively quantify the protein compositions of human MFG and their paired, biochemically isolated membrane fractions to identify proteins specifically enriched on human MFGM. In conjunction with pathway enrichment analyses, our results advance knowledge of the composition and relative quantities of proteins in human and mouse MFG in greater detail, provide a quantitative profile of specifically enriched human MFGM proteins, and identify core cellular systems involved in forming MFGs and MFGMs.

Due to the unique apocrine mechanism of milk lipid secretion, in which portions of the apical plasma membrane, CLD, and apically targeted cellular elements are pinched off from secretory epithelial cells (McManaman 2012, Honvo-Houéto, Henry et al. 2016, Farkaš, Beňo et al. 2020), the protein composition of MFG is predicted to be an aggregate of a select set of proteins captured from the plasma membrane, CLD, and cytosolic fractions of the cell during MFG secretion. Prior studies of human MFG have largely focused on the putative MFGM fractions of these structures (Liao, Alvarado et al. 2011, Yang, Cong et al. 2016), and comparatively few studies have been directed at defining the protein composition of intact MFG and identifying the cellular systems involved in the formation of these structures. Using MS/MS analysis of human MFG proteins separated by 1D SDS-PAGE electrophoresis, Spertino et al (Spertino, Cipriani et al. 2012), identified 13 of the most abundant proteins in human MFG and using LC-MS/MS of iTRAC-labled proteins, Yang et al (Yang, Zheng et al. 2015) identified 520 proteins from human MFG. In contrast, using TIMS/PASEF analysis, we reproducibly quantified relative abundances of 1,824 proteins in intact, freshly isolated, human MFG and 1,587 proteins in mouse MFG, which significantly increases the depth of knowledge about the protein composition of these structures and enhances identification of cellular pathways that contribute to their formation. For intact, freshly isolated human and mouse MFG, we found that the top 20 most abundant proteins in human and mouse MFG were either involved in regulating CLD-membrane interactions during MFG secretion (BTN1A1, XDH/XOR and PLIN2) (Ogg, Weldon et al. 2004, Jeong, Lisinski et al. 2013, Monks, Dzieciatkowska et al. 2016, Monks, Orlicky et al. 2022), were cytoplasmic or cytoplasmic vesicle components (lactadherin (MFGE8), fatty acid binding protein 3 (FABP3) and caseins (casein kappa (CSN3) and casein alpha s1 (CSN1S1) in human, casein beta (Csn2), Csn1s1, casein alpha s2-like A (Csn1s2a) and casein alpha s2-like B (Csn1s2b) in mouse)), or were implicated as regulators of membrane processes and vesicle trafficking (Rab proteins (RAB18, RAB1A), ACTB). Pathway enrichment analysis of human and mouse MFG proteomes also revealed similarities in their corresponding biological pathways. The common, most significantly enriched pathways in MFG from both species were related to vesicular transport, membrane organization, and exocytotic secretion, suggesting that they may represent core cellular processes that contribute MFG formation.

Nevertheless, we also found marked differences between human and mouse MFGs in the relative abundances of specific proteins, which demonstrates differences in their expression and/or incorporation into MFG and in the relative contributions of certain biological processes to MFG formation. For example, lactotransferrin (LTF), which is a secreted protein found in cytosolic vesicles, is 5^th^ in abundance in human MFG versus 147^th^ in mouse; perilipin 3 (PLIN3), which is a CLD-associated protein that is also implicated in endosomal trafficking (Bickel, Tansey et al. 2009), is 7^th^ in abundance in human MFG versus 648^th^ in mouse; and CIDEA, which is a CLD-associated protein in the mouse mammary gland that is implicated as a regulator of milk lipid secretion (Wang, Lv et al. 2012) is 43^rd^ in abundance in human MFG versus 7^th^ for mouse MFG.

We identified proteins specifically associated with human MFGM by comparing the relative abundances of proteins found in intact MFG from individual subjects with their relative abundances in corresponding MFGM fractions in a single LC-MS/MS run. This approach allowed us to directly compare relative protein abundances of MFG and MFGM in individual subjects, which eliminates the potential for inter-run variability and improves the power and quantitative rigor of MFGM protein enrichment analysis over prior studies of human MFGM proteins, which analyzed pooled samples and did not compare protein abundances in MFG and MFGM fractions (Liao, Alvarado et al. 2011, Yang, Zheng et al. 2015, Yang, Cong et al. 2016, Zhang, Jiang et al. 2021). Using differences in relative abundances between MFG and isolated MFGM, we found 628 proteins whose relative abundances were significantly increased MFGM after correcting for multiple comparisons. The top 20 MFGM-enriched proteins included those demonstrated to contribute to CLD docking in mouse models of milk lipid secretion - BTN1A1 (#1), XDH/XOR (#2), PLIN2 (#3) and CIDEA (#6) (Monks, Dzieciatkowska et al. 2016, Monks, Orlicky et al. 2022) which have previously been reported to be major MFGM proteins (Reinhardt and Lippolis 2008, Thum, Wall et al. 2022). We also found several known membrane proteins among the top 20 enriched proteins, including the ATP-binding cassette transporter G member 2 (ABCG2, #7) that is linked to riboflavin transport in human milk (Golan and Assaraf 2020), cytochrome B5 type A (CYB5; #9), a redox enzyme previously found on MFGM in humans and other species (Jarasch, Bruder et al. 1977), CD59 molecule (CD59 blood group) (CD59, #10), a complement factor previously identified in human MFG (Hakulinen and Meri 1995), toll-like receptor 2 (TLR2, #11), which has been detected on bovine MFGM (Reinhardt and Lippolis 2008); and calcineurin-like EF-hand protein (CHP1, #17), a regulator of endocytosis (Janzen, Mendoza-Ferreira et al. 2018) and calcium and integrin binding 1 (CIB1, #19), a suppressor of integrin activation (Kim, Ye et al. 2011), which have not been reported to be MFGM proteins previously. In addition, lipid synthesis enzymes acetyl CoA synthetase long chain family members 1 and 4 (ACSL1 #12, ACSL4 #13) and lanosterol synthase (LSS, #16) are among the top human MFGM-enriched proteins, ACSL1 and LSS have been detected previously on bovine MFGM (Reinhardt and Lippolis 2008). Other abundant human MFGM-enriched proteins that have been detected on MFGM previously are lactalbumin alpha (LALBA, #4) which regulates lactose synthesis and is reported to have antimicrobial properties (Charlwood, Hanrahan et al. 2002); eukaryotic translation elongation factor 1 alpha 1 (EEF1A1, #5), an actin-binding protein that contributes to the regulation of epithelial cell junctions (Erasmus, Bruche et al. 2016); calcium binding proteins calnexin (CANX, #15) and S100 calcium binding protein A1(S100A1, #8) (Reinhardt and Lippolis 2008, Yang, Cong et al. 2016); and syntaxin binding protein 2 (STXBP2, #20) (Reinhardt and Lippolis 2008). We did not find that 2’,3’-cyclic nucleotide 3’-phosphodiesterase (CNP, #14), or Ribophorin 1 (RPN1, #18) had been detected on MFGM previously.

By PCA, we found differences in relative abundances of human MFGM-enriched proteins between transitional vs mature lactation timepoints that was driven primarily by the significantly increased abundance of BTN1A1 in MFGM over time. Although previous human and bovine studies have reported that the expression of protein mediators of CLD-membrane interactions including BTN1A1, XDH/XOR, PLIN2 and CIDEA increase between colostral and mature phases of lactation (Reinhardt and Lippolis 2008, Yang, Cong et al. 2016), we did not detect significant differences in the relative abundances of XDH/XOR, PLIN2 or CIDEA in MFGM fractions from transitional and mature milk in our cohort, which suggests that molecular complexes involved in docking CLDs to the apical membrane are largely established in humans by the 2^nd^ week of lactation.

Pathway analysis of the proteins enriched in human MFGM identified lipid biosynthesis, localization, and metabolism as among the most significantly enriched pathways. In addition, several significantly enriched pathways related to MFG proteins were also found to be significantly enriched in analysis of MFGM-enriched proteins. The enriched pathways include vesicle transport, membrane organization and exocytosis, which is consistent with the proposal that vesicular compartments contribute to MFGM formation (Wooding 1973). In addition, there is significant enrichment of multiple ER-associated pathways in human MFGM-enriched proteins, including N-linked glycosylation, response to stress, and protein processing. The enrichment of these pathways in the human MFGM is consistent with studies in mice proposing that ER proteins contribute to MFGM formation (Wu, Howell et al. 2000, Honvo-Houéto, Henry et al. 2016). Each of the significantly enriched ER pathways is related to protein folding, which suggests that discrete ER elements required for correct protein folding and/or protein quality control may be specifically directed to apocrine lipid secretion. Consistent with this mechanism, our proteomics data show that several proteins implicated in tethering the ER to the plasma membrane (Li, Qian et al. 2021) are enriched in isolated human MFGM, including VAMP associated protein A (VAPA, #76), extended synaptotagmin proteins ESYT1 (# 207) and ESYT2 (# 337) and oxysterol-binding protein like proteins – OSBPL2 (# 239), OSBPL1A (# 364) and OSBPL8 (# 423).

Importantly, we found that the relative abundances of several proteins reported to be major MFGM proteins from humans or other species (Thum, Wall et al. 2022) either did not differ significantly between MFG and MFGM fractions or were significantly more abundant in the MFG fraction. The relative abundance of lactadherin (MFGE8), for instance, was significantly greater in human MFG compared to corresponding MFGM fractions, whereas relative abundances of FABP3, mucin1 (MUC1); and CD36 molecule (CD36) did not differ significantly between human MFG and MFGM. These results provide evidence that for some proteins previously thought to be enriched on MFGMs, their membrane association may be comparatively weak or their abundances in other MFG compartments may be comparable to their membrane abundances.

We found 205 proteins with significantly greater abundances in MFG compared to their corresponding MFGM factions. In addition to MFGE8, LTF and CSN3, which are found in secretory vesicles, we found significant increases in the abundances of several cytosolic proteins including enolase 1 (ENO1), carbonic anhydrase-6 (CA6), glyceraldehyde-3-phophate dehydrogenase (GAPDH) as well as numerous ribosomal proteins (RPS17, RPS7, RPLP2, RPS10, RPL38, RPS15 and RPS15A) in MFG relative to MFGM fractions. Collectively these data are consistent with the apocrine mechanism of lipid secretion, which is proposed to capture soluble and vesicular fractions of the cytosol in addition to CLD. However, mouse data have shown that apocrine lipid secretion is facilitated by, but does not require, contact between CLD and the apical plasma membrane (Monks, Dzieciatkowska et al. 2016, Monks, Orlicky et al. 2022). Thus, variations in the extent of CLD docking or in maternal physiological processes provide opportunities for variable combinations of cellular membranes and cytoplasmic components in secreted MFG.

In summary, we have used ultra-sensitive TIMS/PASEF proteomics to identify and define relative abundances of proteins in human MFG and MFGM and mouse MFG in greater detail. Coupled with bioinformatic pathway analyses, our results provide new information about the protein compositions of human and mouse MFGs and the cellular processes that contribute to their formation. By comparing relative abundances of human MFG and MFGM proteins we were able to identify a set of proteins that are specifically enriched on human MFGM. Collectively these data provided new insight into the protein compositions of human MFG and MFGM and the cellular processes involved in their formation, which we speculate will help to define the importance of these unique structures in infant nutrition.

## Supplemental Figures

**Supplemental Figure 1.**
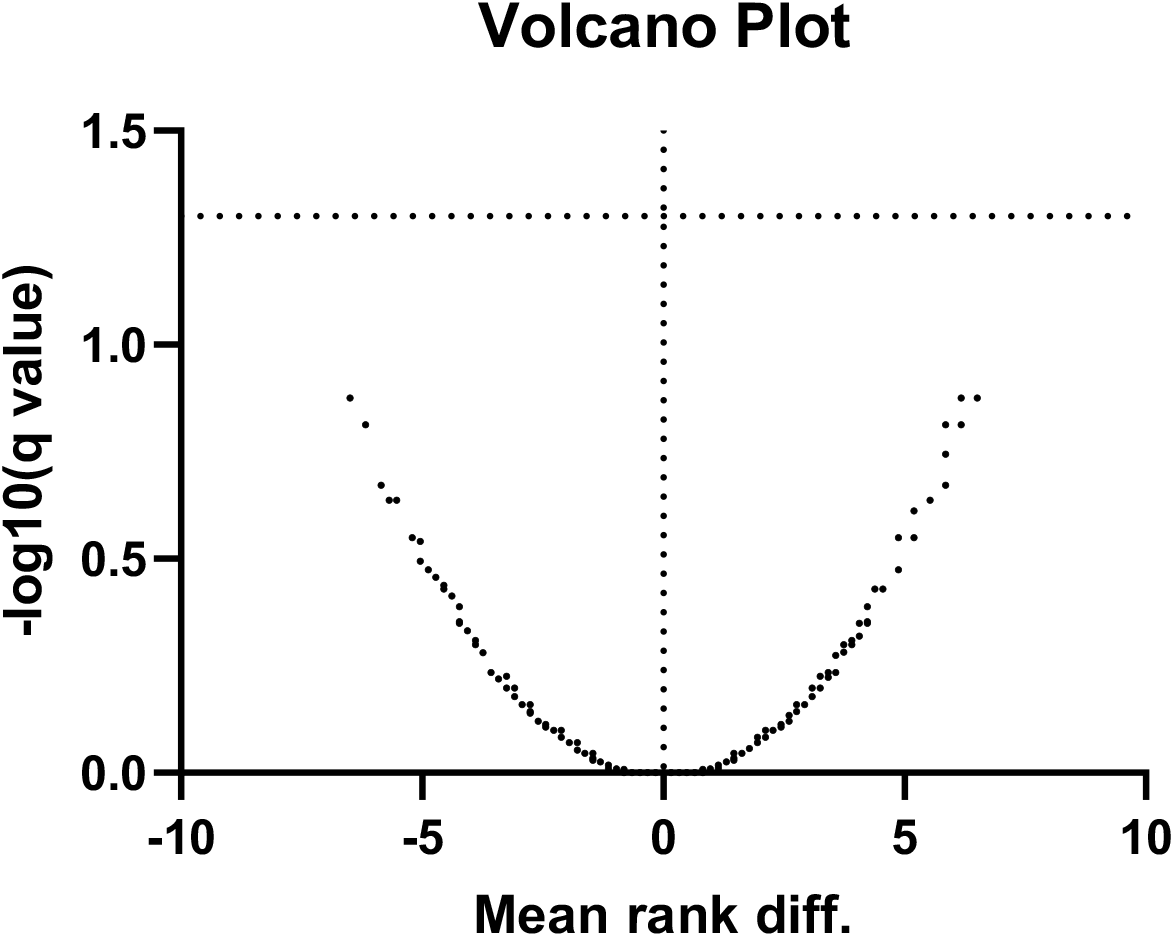
Multiple non-parametric (Mann-Whitney) unpaired t-tests of MFGM-associated proteins between early (2wk) and mature (2-5mo) lactation timepoints with FDR correction for multiple comparisons (5%).

**Supplemental Figure 2.**
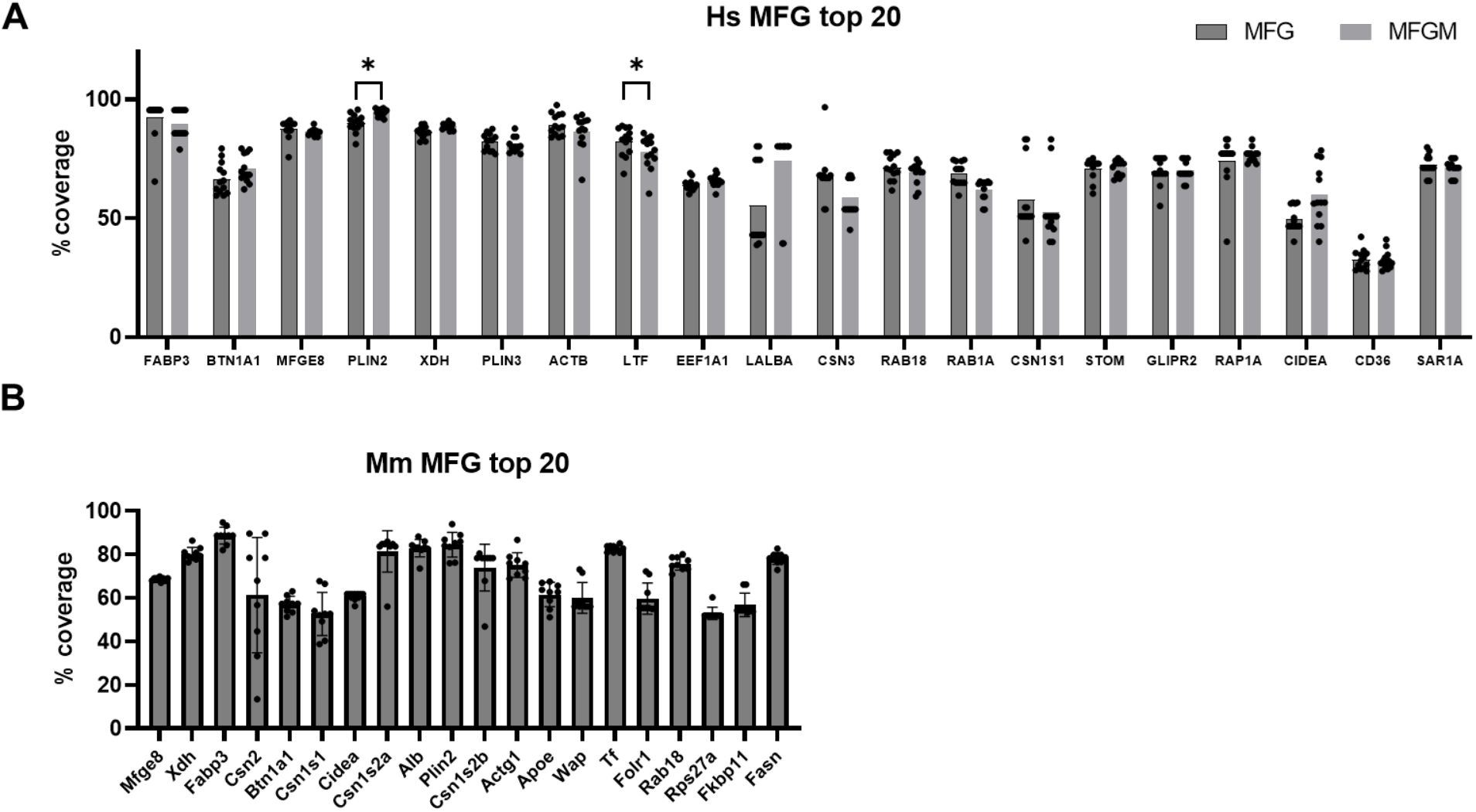
Percent coverage of top 20 human MFG-(dark gray bars) and MFGM-associated (light gray bars) proteins (**A**). Multiple paired t-tests, adjusted for multiple testing with the Holm-Šídák method. *p<0.05. Percent coverage of top 20 CD1 mouse MFG-associated proteins (**B**).

**Supplemental Figure 3.**
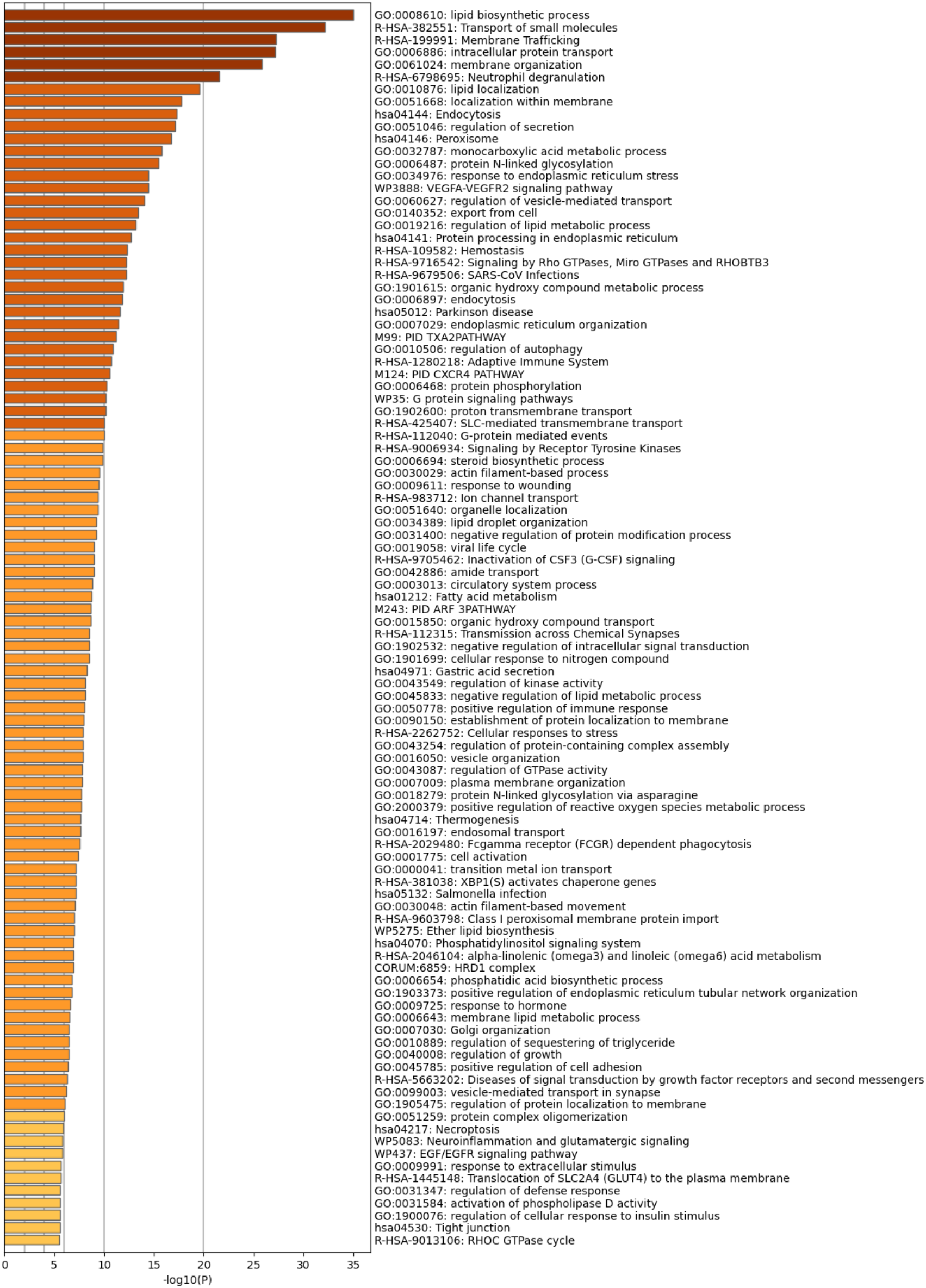
Metascape pathway enrichment analysis of human MFGM-enriched proteins, calculated by difference.

**Supplemental Figure 4.**
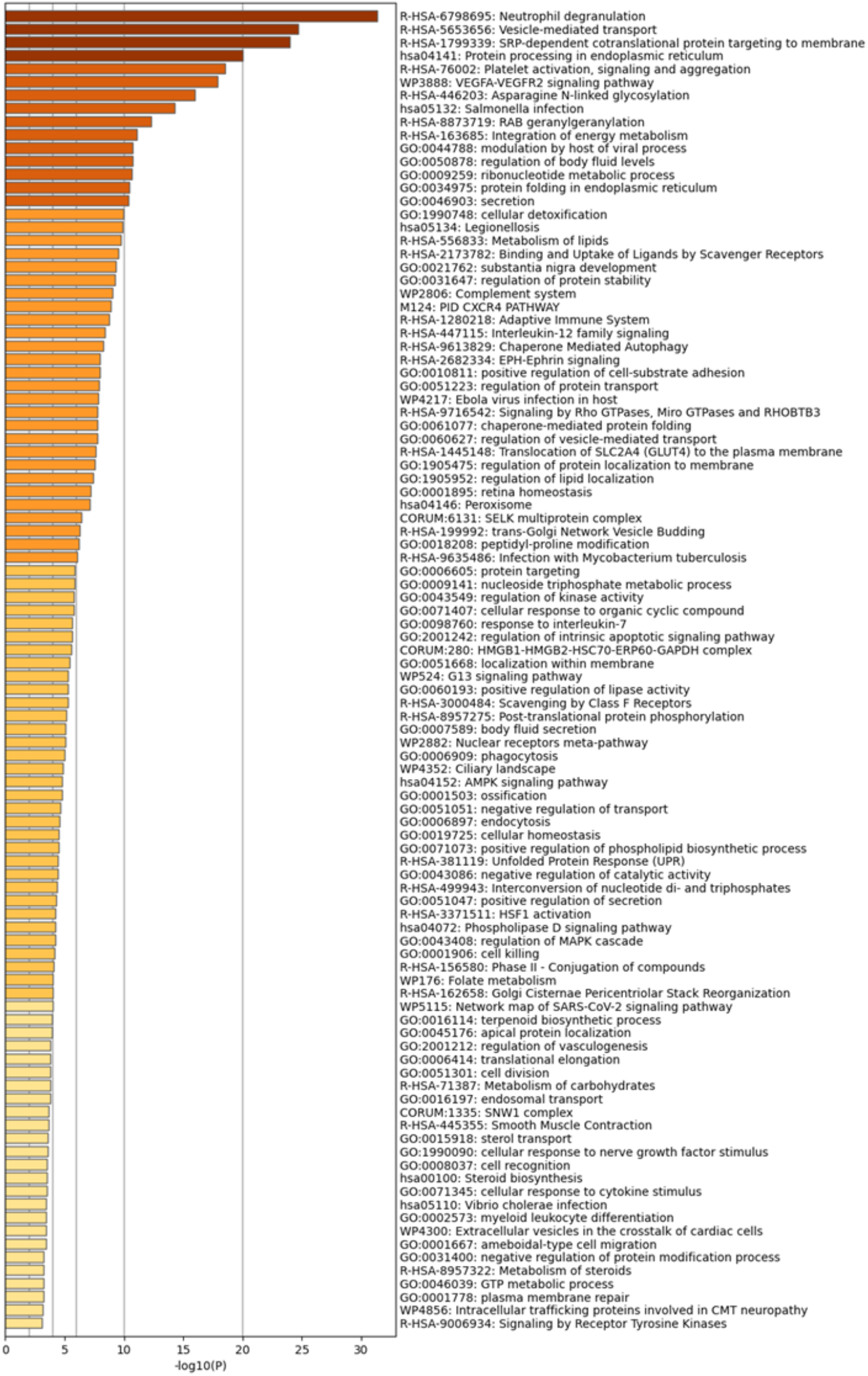
Metascape pathway enrichment analysis of the top 200 (sorted by median) human MFGM-associated proteins.

**Supplemental Figure 5.**
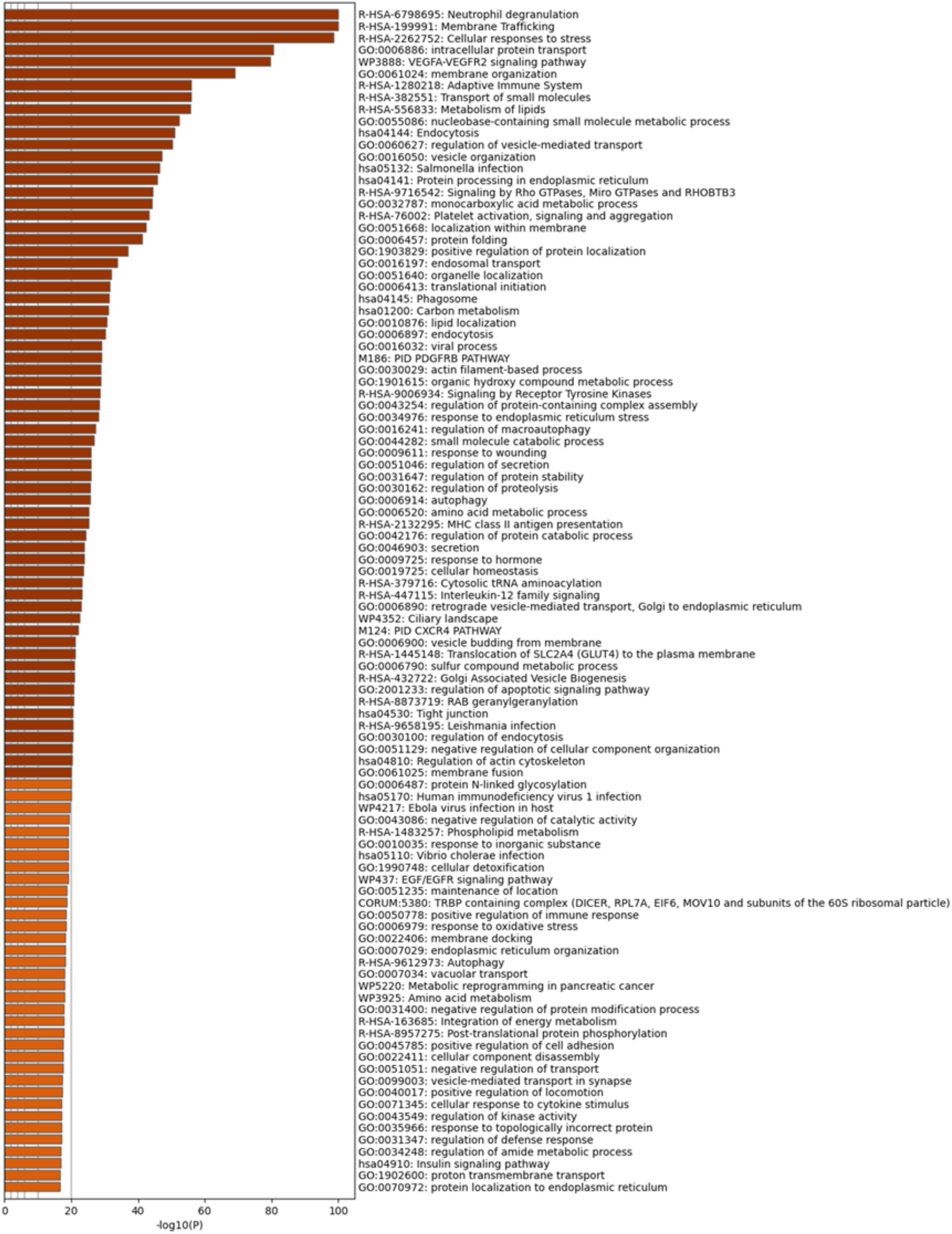
Metascape pathway enrichment analysis of all human MFGM-associated proteins.

**Supplemental Figure 6.**
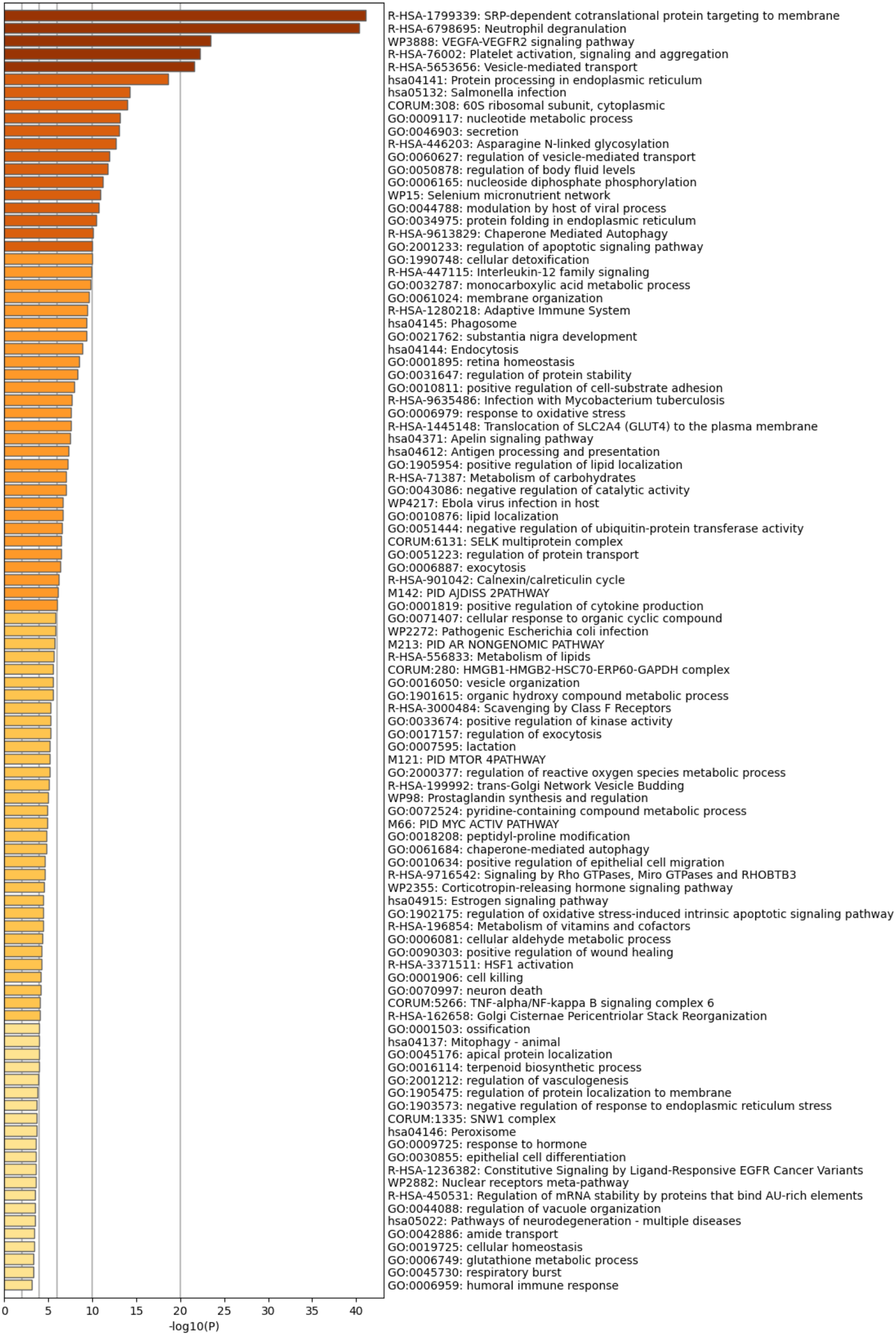
Metascape pathway enrichment analysis of the top 200 (sorted by median) human MFG-associated proteins.

**Supplemental Figure 7.**
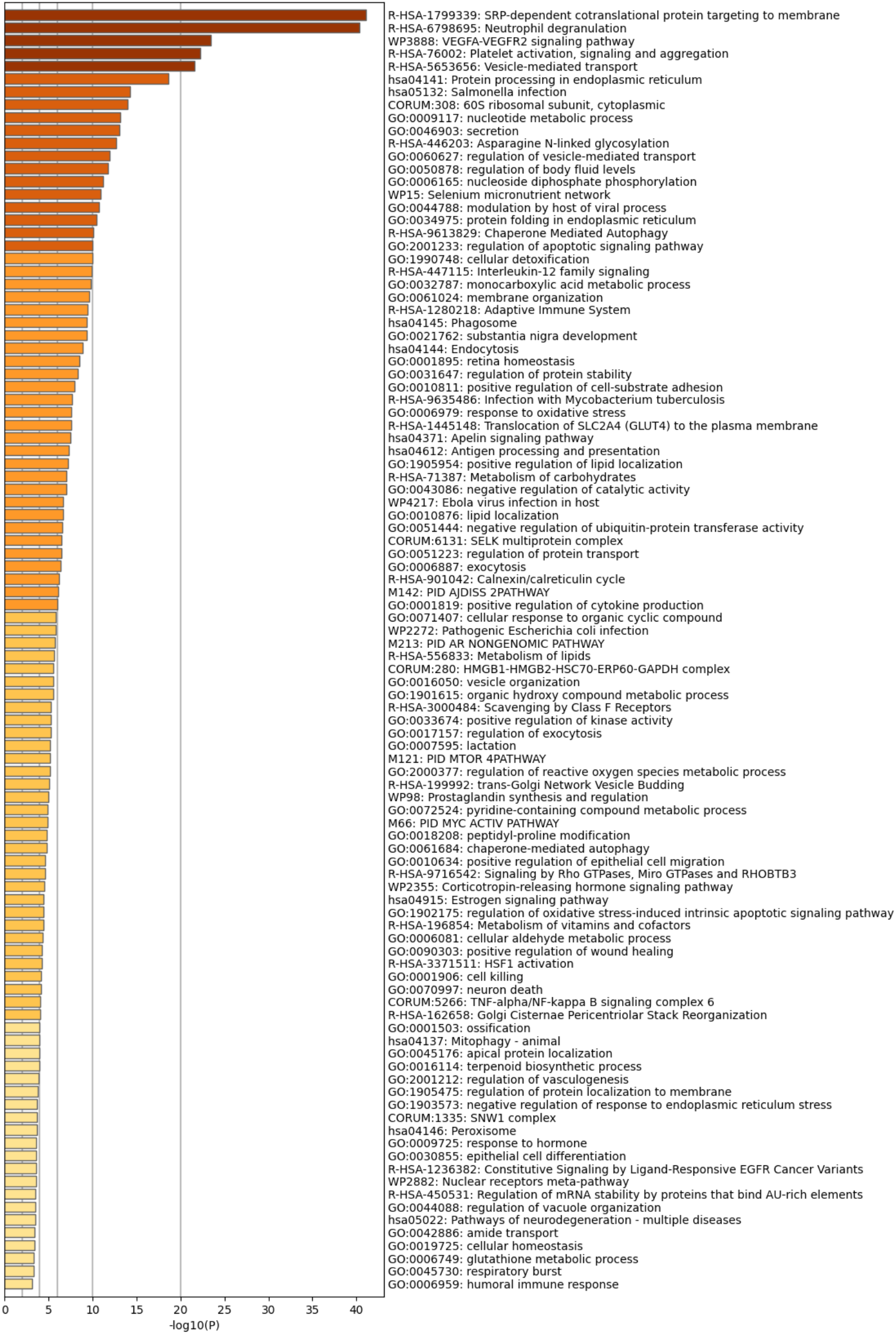
Metascape pathway enrichment analysis of all human MFG-associated proteins.

**Supplemental Figure 8.**
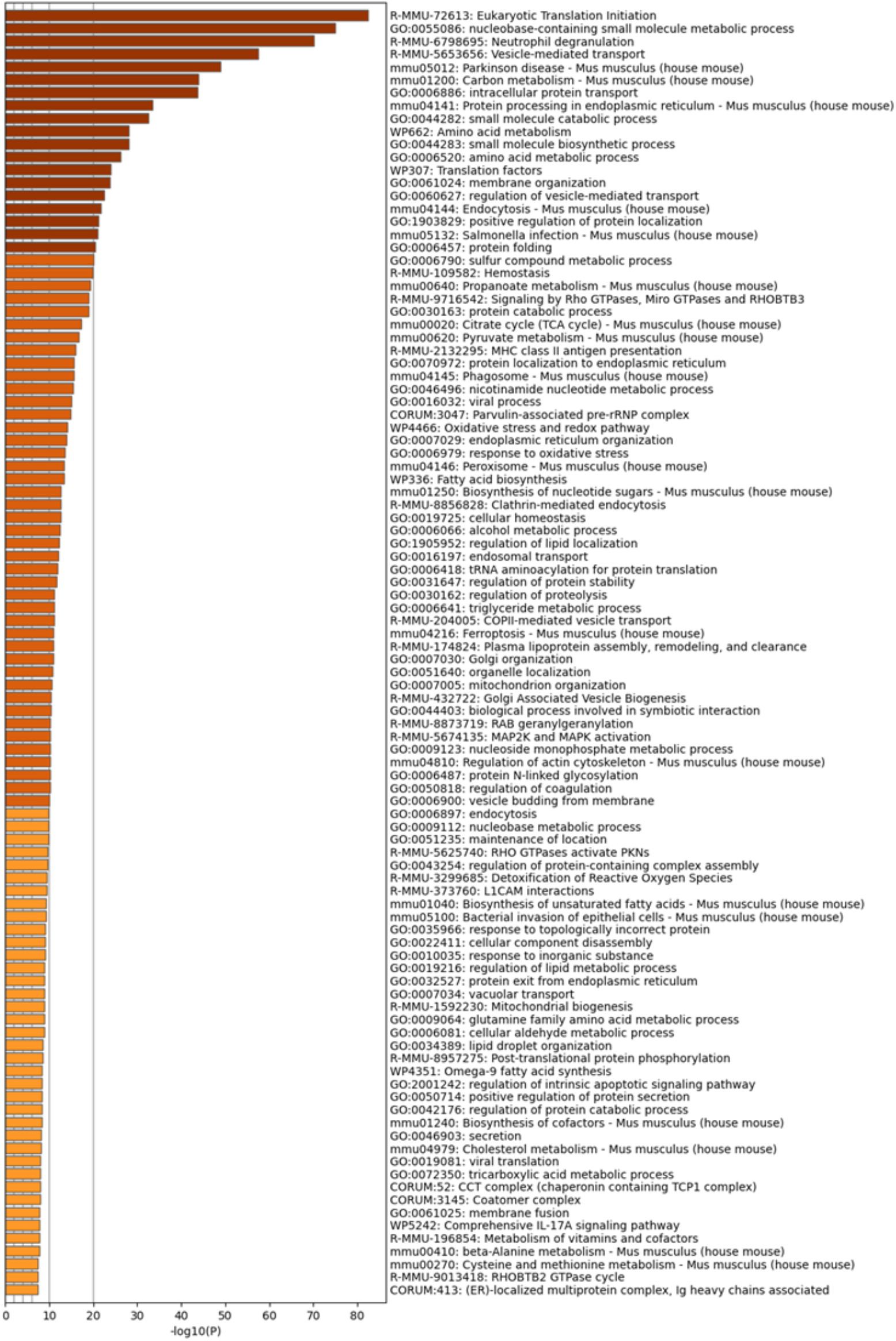
Metascape pathway enrichment analyses of mouse MFG-associated proteins.

**Supplemental Figure 9.**
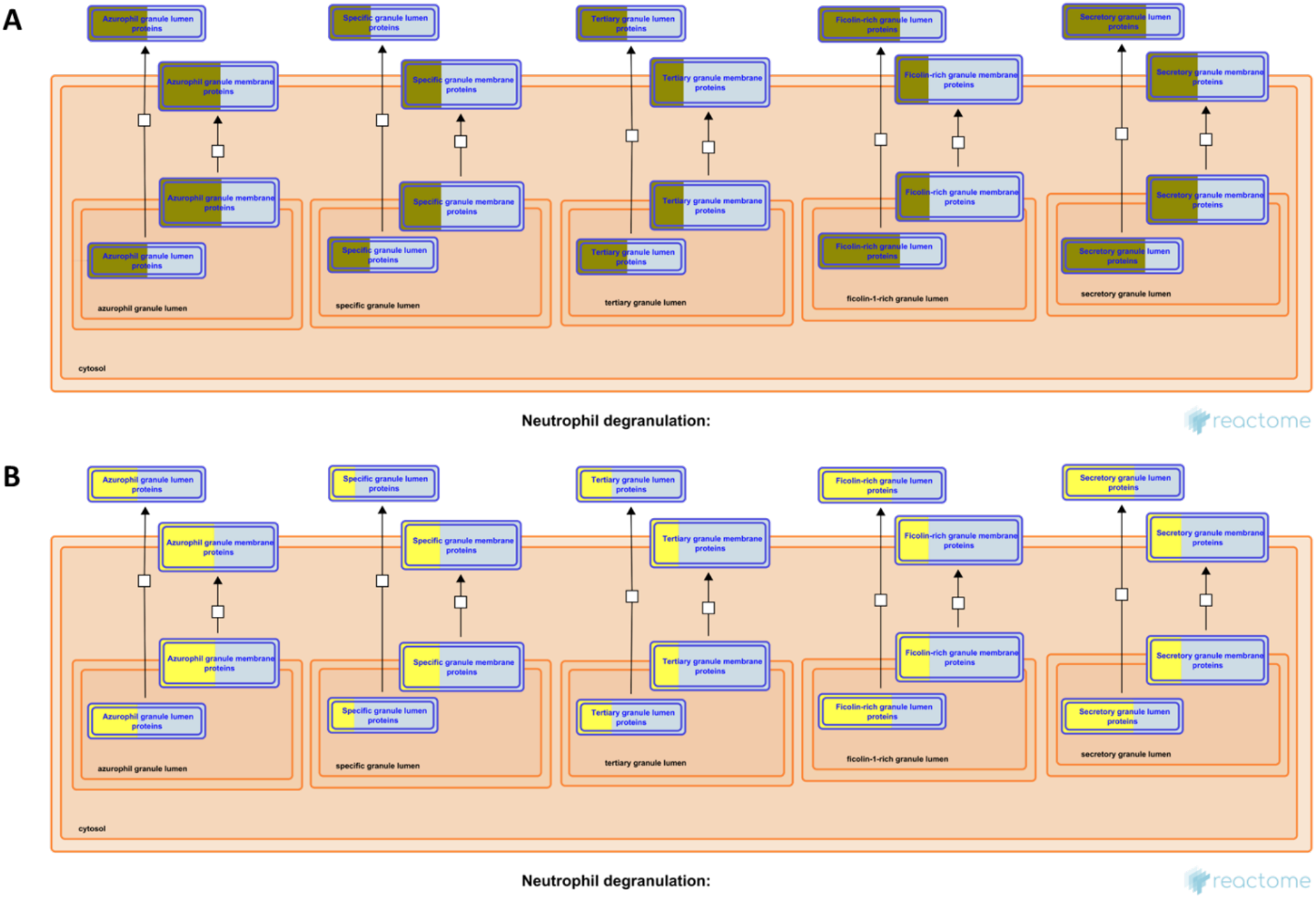
Reactome pathway diagrams with overlaid overrepresentation pathway enrichment analyses of human (**A**) and mouse (**B**) MFG-associated proteins in the neutrophil degranulation pathway. Entities are re-colored (gold for human, yellow for mouse) if they were represented in the submitted data set.

**Supplemental Figure 10.**
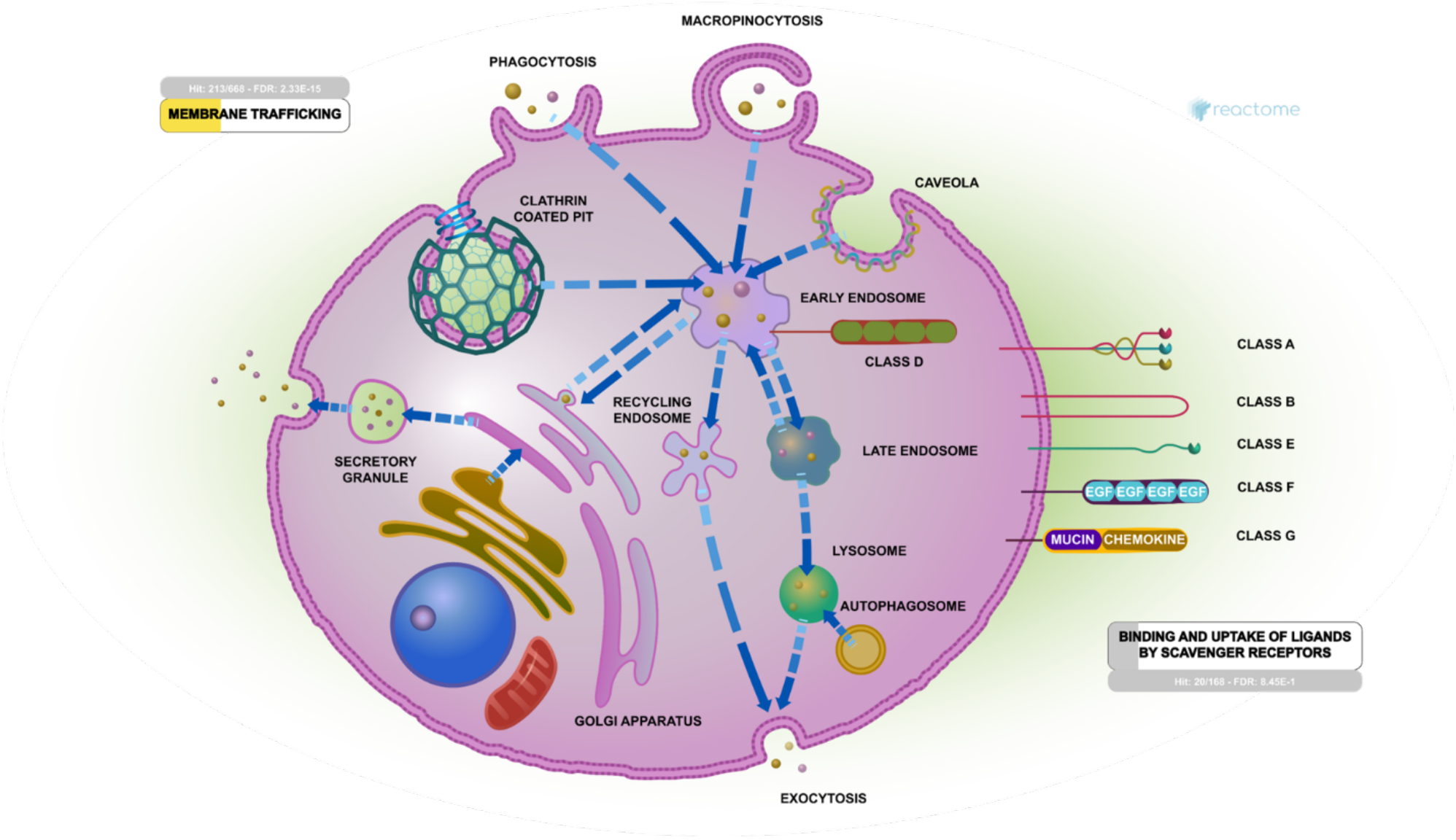
Reactome pathway diagrams with overlaid overrepresentation pathway enrichment analyses of human MFG-associated proteins in the vesicle mediated transport pathway. Entities are re-colored (yellow if statistically significant, gray if not statistically significant) if they were represented in the submitted data set.

**Supplemental Figure 11.**
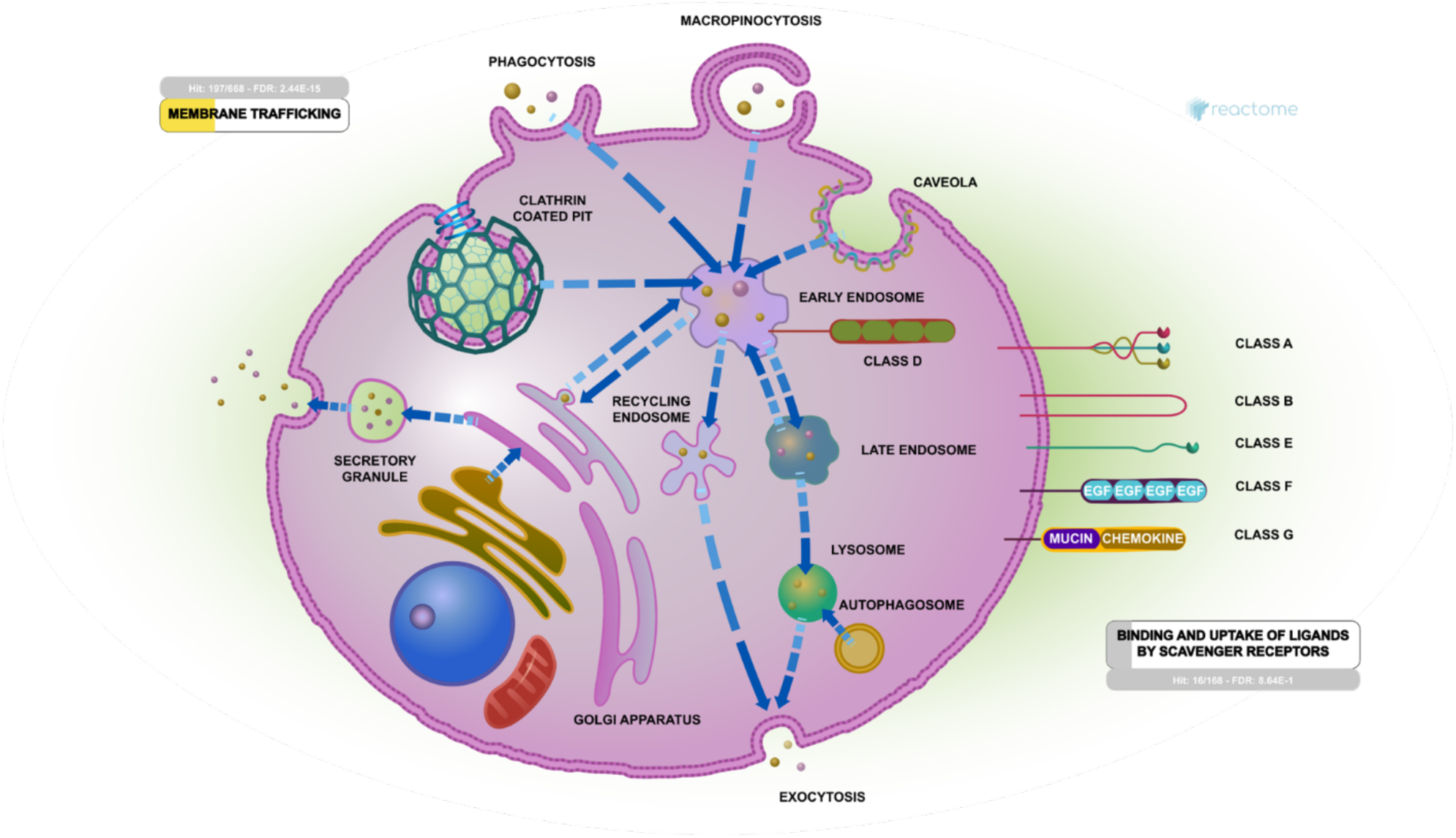
Reactome pathway diagrams with overlaid overrepresentation pathway enrichment analyses of mouse MFG-associated proteins in the vesicle mediated transport pathway. Entities are re-colored (yellow if statistically significant, gray if not statistically significant) if they were represented in the submitted data set.

